# Active Perception during Angiogenesis: Filopodia speed up Notch selection of tip cells *in silico* and *in vivo*

**DOI:** 10.1101/2020.08.22.261263

**Authors:** Bahti Zakirov, Georgios Charalambous, Irene M. Aspalter, Kelvin Van-Vuuren, Thomas Mead, Kyle Harrington, Raphael Thuret, Erzsébet Ravasz Regan, Shane Paul Herbert, Katie Bentley

**Affiliations:** Cellular Adaptive Behaviour Lab, Francis Crick Institute, London, United Kingdom; Department of Informatics, King’s College London, London, United Kingdom; Division of Developmental Biology and Medicine, University of Manchester, Manchester, United Kingdom; Virtual Technology and Design, University of Idaho, Moscow, Idaho, USA; Center for Vascular Biology Research, Beth Israel Deaconess Medical Center, Department of Pathology, Harvard Medical School, Boston, USA; Department of Biology, The College of Wooster, Wooster, Ohio, USA

**Keywords:** Angiogenesis, Active Perception, Basal Cognition, Morphogenesis, Notch Lateral Inhibition, Filopodia

## Abstract

How do cells make efficient collective decisions during tissue morphogenesis? Humans and other organisms utilize feedback between movement and sensing known as ‘sensorimotor coordination’ or ‘active perception’ to inform behaviour, but active perception has not before been investigated at a cellular level within organs. Here we provide the first proof of concept *in silico/in vivo* study demonstrating that filopodia (actin-rich, dynamic, finger like cell-membrane protrusions) play an unexpected role in speeding up collective endothelial decisions during the time-constrained process of ‘tip cell’ selection during blood vessel formation (angiogenesis).

We first validate simulation predictions *in vivo* with live imaging of zebrafish intersegmental vessel growth. Further simulation studies then indicate the effect is due to the coupled positive feedback between movement and sensing on filopodia conferring a bistable switch-like property to Notch lateral inhibition, ensuring tip selection is a rapid and robust process. We then employ measures from computational neuroscience to assess whether filopodia function as a primitive (‘basal’) form of active perception and find evidence in support. By viewing cell behaviour in tissues through the ‘basal cognitive lens’ we acquire a fresh perspective on not only the well-studied tip cell selection process, revealing a hidden, yet vital, time-keeping role for filopodia, but on how to interpret and understand cell behaviour in general, opening up a myriad of new and exciting research directions.

## Introduction

Morphogenesis of multicellular tissues and organs is a remarkably precise, robust, and adaptive process[1] [2, 3]. How do cells manage to coordinate their behaviour so precisely and rapidly, given the complexities and time constraints of the developing embryo environment they reside within? We will argue, and provide the first proof of concept evidence, that cells in tissue development have evolved a basal cognitive process of active perception in order to deliver on the adaptive problem-solving required of them during morphogenesis.

We focus here on angiogenesis, the process of new blood vessel sprouting from pre-existing ones. Angiogenesis is critical for tissue development, homeostasis, and repair across organs in the body and is frequently dysregulated in disease[4-6]. Angiogenesis is driven first and foremost by sprouting endothelial cells (ECs) as they form the new blood vessel walls. EC behaviour is highly dynamic and needs to be tightly coordinated in space and time to effectively perfuse the surrounding tissue. Despite extensive progress to better understand angiogenesis since the discovery of its role in cancer progression in the 1970s[7, 8], many aspects of the timing and cell dynamics involved are not yet understood, largely due to an early lack of live-imaging methods [9]. Vascular networks are highly specialized morphologically and functionally in different organs of the body [10, 11]. Although a remarkably consistent core set of molecular pathways and processes have been identified that appear to lie at the heart of angiogenesis across tissues and species [12] organ-specific fine-tuned adaptations to this core are beginning to be uncovered ([13, 14]. It is clear that much remains to be understood about how ECs gain information about the local tissue in order to adapt and construct organ-specific vascular network topologies and functions[15] [16, 17].

In the first core step of angiogenesis, ECs in a vessel collectively decide (‘select’) who will be the most migratory (the ‘tip’ cell) and lead a new vessel ‘sprout’ that branches out from the pre-existing vessel (Fig. 1a) the other cells then follow behind (‘stalk cells’). Tip cells are well-studied and characterized by their long thin, highly dynamic protrusions - filopodia (Fig.1b). Two sprouts eventually fuse to form vessel loops that can then support blood flow (Fig.1a). This process repeats iteratively building up a dense looped network (the ‘primary plexus’) that is later remodelled and pruned through cell rearrangement and blood flow to a more familiar hierarchical tree like structure[18] [19].

**Figure 1.**
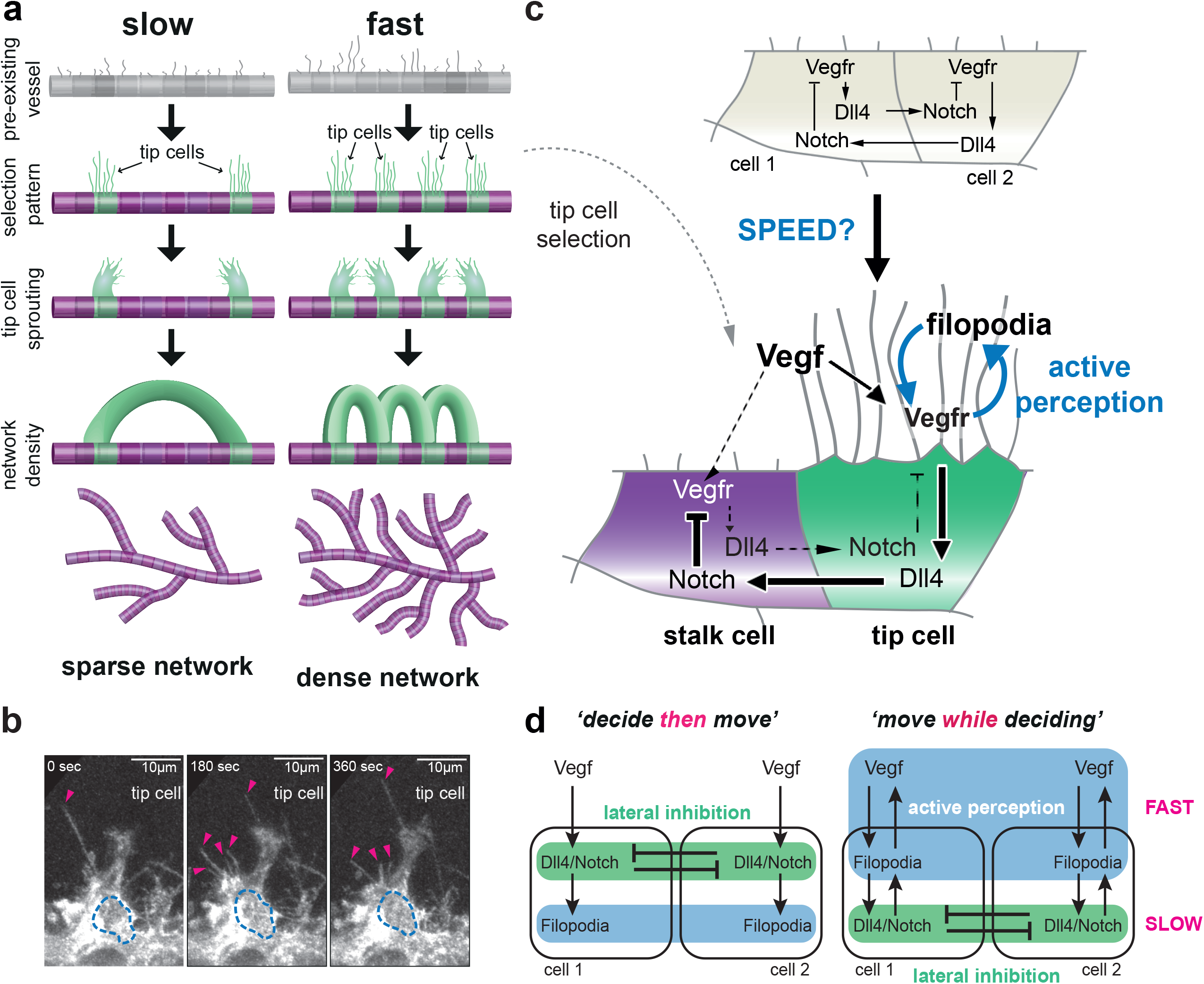
Mechanisms of tip selection during vascular network formation (angiogenesis): **a)**A pre-existing vessel becomes activated by factors in a hypoxic tissue environment (low in oxygen) starting the process of tip selection where an alternating “salt and pepper” pattern of tip (green) and stalk cells (purple) emerges. Tip cells then migrate out, extending new branch “sprouts”, which fuse (anastomosis) to form loops supporting blood flow. The speed of selection determines the branch spacing properties of the resulting blood vessel network. Fast patterning leads to a denser network, slow patterning leads to a sparser network.**b)**Representative time-lapse images of filopodia extension from a tip cell migrating from the dorsal aorta (DA) into upwards to form the intersegmental vessel (ISV) of a Tg(kdrl:ras-mCherry)s896 embryo. The dotted blue lines highlight the nucleus of the tip cell. Pink arrows mark filopodia. **c)**The vegf /notch tip selection pathways. Vegf interacts with Vegfr-2, expressed at the surface of the endothelial cells, and on filopodia. Vegfr-2 activation triggers Dll4 gene expression up-regulation and actin polymerisation causing filopodia extension (e.g. via Pi3K). In turn, Dll4 ligands activate Notch receptors on the neighbour cell triggering cleavage of the Notch intracellular domain (NICD), which translocates to the nucleus reducing gene expression of Vegfr-2 receptors (and other pathways). Differences in Vegfr activation are amplified by this lateral inhibition competition between the cells until one cell has more Vegfr-2 receptors (which can continue stimulating the actin migratory response) and Dll4 (tip cell) inhibiting its neighbour (stalk cell, which has higher Notch activation levels and less Vegfr). **d)**Two ways to conceptually frame how notch decisions take place. One way corresponds to the prevailing view of the process-“Decide then Move” - which designates filopodia as the end result of the selection decision-making process – prominent on migrating tip cells. The “Move While Deciding” view, which takes into account the relative timescales of events, implicates filopodia ahead of Notch, driving the decision-making process.

Crucially, we recently found through an *in silico* / *in vivo* study that if the cells are slower to collectively select the new tip cells, then this has real consequences for the resulting vascular network topology: the slower the selection, the less dense the network (Fig 1a) [17]. We have begun to uncover a growing number of ‘temporal regulators’ of EC collective decisions (e.g. tissue-derived factors like Semaphorins[17] and cell-specific factors such as the tetraspanin-like protein Tm4sf18 [20]) that can fine tune branch density in a potentially organ-specific manner.

Here we demonstrate that filopodia, rather than just being features of the *selected* migratory tip cells, provide a vital, overlooked ‘temporal regulator’ role, keeping the selection ‘decision-making’ process timely. This highlights a novel role for cell shape changes in enhancing cell decision-making. Namely, filopodia extensions and retractions (*movements*) alter receptor locations and activations (*sensations*), amplifying differences between the cells in a positive feedback cycle. This is a form of sensorimotor coordination which, we argue, could constitute a basal form of Active Perception (AP). To our knowledge, Active Perception has until now only been studied and identified in higher organisms and prokaryotic cells [21, 22].

First, we will give a brief introduction and context to the biological and basal cognitive concepts proposed in this study. We detail the tip selection process and present our time-ordering argument for the overlooked role of filopodia during selection. Second, we present our original research findings providing: a) the first *in vivo* evidence during zebrafish embryo development that filopodia loss slows selection as predicted *in silico* in our cell-based simulations with the previously well-validated MemAgent-Spring model (MSM) [23-27]b) simulations identifying that filopodia enable rapid and robust selection by conferring bistable decision-making properties to individual ECs due precisely to the feedback coupling of sensation and movement of filopodia and c) a simple information theory metric to extract quantitative ‘signatures’ of active perception, which has potential to be extended and applied to future *in vitro* / *in vivo* and *in silico* experiments aiming to detect and further delineate the role of cell level AP in tissues.

Filopodia are well-documented during angiogenesis in most organs in the body. We propose that their time-keeping role during selection could be a ubiquitous part of the angiogenic process. Moreover, AP properties could be endowed to other types of cells by different processes and shape-changing mechanisms. The scope of this study, however, is to provide the first proof of concept and simple, small step. We therefore conclude with a discussion into the potentially wide-reaching new research avenues this study opens up, both drilling down into the several different detailed molecular mechanisms potentially underlying AP in cells of different tissue-beds and ways that AP could present a novel ‘higher-level’ target to therapeutically modify cell behaviour to tune branching to a desired density in diseased or engineered tissues.

## Background

### Notch Tip Cell Selection is too slow

The process to select new tip cells is currently considered to be well-understood and already well-characterised to occur via Notch-Dll4 lateral inhibition [28-30] triggered primarily by Vascular Endothelial Growth Factor (Vegf) released when the surrounding tissue becomes hypoxic (low in oxygen) (Fig. 1c) [31, 32]. However, this widely accepted selection mechanism does not consider timing or the specific dynamics of the process. We recently argued that this has caused us to miss a critical, speed-controlling feature, required for normal selection [9, 33]. The *multiple* cycles of gene expression required by Notch-Dll4 lateral inhibition to amplify initially small differences in input Vegf signal of neighbouring ECs on a vessel to select the heterogenous pattern of tip cells is seemingly incompatible with the rapid, dynamic changes in EC state, identity, and behaviour observed in angiogenesis[34, 35]. We recently showed that selection of tip cells is a specifically time constrained step of the process during zebrafish intersegmental vessel formation [20]. In this phase of zebrafish development, tip cells emerge within a ‘selection window’ of approximately 8 hrs ending at 24 hrs post fertilization (hpf), after which no new tip cells are selected. One cycle of Notch-Dll4 lateral inhibition takes approximately 4-6 hours *in vitro* in mouse ECs [36] suggesting the currently accepted selection process would have to take closer to 16-24hrs to select each tip cell. This is true even accounting for the more rapid gene expression cycles in zebrafish (approximately 2 hours for one cycle, [37]. Thus, we hypothesize that Notch lateral inhibition alone is too slow alone to account for *in vivo* selection.

### A role for filopodia as time-keepers during collective cell decisions

So how do endothelial cells make rapid tip selection decisions? Early simulations with the MSM model of Vegf-Notch selection first pointed to filopodia as an unexpected temporal regulator of the process [24], however the significance of those early simulations was not clear until now. Given that filopodia are normally used as a marker of the selected tip cell phenotype, this was a surprising prediction - how could the consequence of selection be part of the cause? We recently presented a carefully revised time-ordering of the precise events during selection, which highlighted a missed role for filopodia *before* tip cells are selected [9].

#### Time-ordered account of selection

When Vegf binds and activates Vegf receptors (Vegfr) on existing endothelial cells, the cells respond on the *order of seconds* by locally polymerizing actin and pushing out the cell membrane into thin protrusive shapes – filopodia - which are then rapidly extended and retracted [38] [39]. These in turn move the Vegf sensors through the environment, as they reside on the deforming cell membrane. This creates a rapid positive feedback to Vegf signalling, which in itself can amplify and structure the differential Vegf input between neighbouring cells, requiring fewer cycles through the slow *(on the order of hours)* Notch gene-regulatory lateral inhibition mechanism to ‘stabilise’ the decision. Thus, filopodia are chronologically primary in the decision-making process, and Notch is secondary – cells “move while deciding” (Fig 1c), contrary to the prevailing view that filopodia movement are associated with the selected tip cells “decide then move” [40-42].

The role of filopodia in angiogenesis has been surprisingly elusive, despite their clear abundance during the process. They were first identified as a guidance mechanism in angiogenesis, helping the selected tip cells, to follow Vegf gradients [43]. However, they were later deemed to be ‘dispensable for guidance’ in zebrafish ISV sprouting, albeit still slowing down sprouting when reduced [44]. Filopodia together with lamellipodia are thought to anchor tip cells to extracellular substrates by focal contact points, together pulling the tip cell forward during migration and sprouting [45] though filopodia are known to be inhibitory to migration in other cell migration contexts [46]. They play a key role in facilitating the formation of anastomoses with other ECs [25, 47, 48]. However, these filopodia likely play a number of roles on ECs, both before and after tip cell phenotypes are selected. Filopodia have previously been defined to expand the spatial limits of cell-cell communication during Notch lateral inhibition patterning in drosophila bristle selection [49], by allowing them to reach out and contact more distant cells, but have not to our knowledge before been investigated as a temporal adaptor of Notch patterning, altering the speed and dynamic properties of Notch lateral inhibition.

### Filopodia and Active Perception

The chronological ordering of the selection process (Fig. 1d) brings movement to the forefront and highlights a potential role for ‘sensorimotor’ feedback in informing Notch selection decisions. ‘Moving while deciding’ and sensorimotor feedback are central to early definitions of Active Perception (AP) in the field of psychology[50-53] where AP is well known to enhance human decision-making and execution of other cognitive tasks such as language acquisition [54]. Thus, we consider whether filopodia may constitute a simple cell level form of AP enhancing collective tip selection decisions.

A particularly relevant example of the necessity of active perception at the human scale is found in the study of saccadic eye movements. In this context, sensory input obtained by a stationary /passive individual is not sufficient to facilitate its behaviour without local movements of sensors on the otherwise immobilized body. The necessity of movement is related to the anatomy of a human eye-which has a highly concentrated cluster of sensory cells at a region known as the fovea. The area of the fovea is small, and sensory acuity falls off rapidly in regions further away from it. Vision is largely facilitated by the eye undergoing abrupt movements, purposefully darting the fovea around visual scenes and gathering relevant information (at a rate comparable to an optimal Bayesian model). An observer would not be able to accurately characterize a visual scene without the ability to move their fovea. [55].

Indeed, biology across scales is rife with examples where AP is required for behaviour including organisms ranging from E Coli to C Elegans, Drosophila, and Humans. [21, 55-57]. At the closest scale to ECs - single prokaryotic cells, the run and tumble behaviour of E Coli during chemotaxis is a well-established example of such a case, where the noise induced by the random arrival of chemoattractant molecules makes it physically impossible for a stationary bacterium to know which direction is “up” the chemoattractant gradient ([56, 58-60]. Though their particular modes of movement may differ, the commonality between these examples is that the agents in question all must utilize coordinated sensorimotor activity to gather enough information to facilitate their behaviours.

We aim here, however not just to make a simple analogy that filopodia on ECs are akin to active perception in higher or single-celled organisms, but to take the first steps in creating the analysis tools needed to rigorously prove that the effect of filopodia on EC selection speed is specifically due to the coupling of movement and sensation that they generate. As AP has become more widely recognised and studied in a diversity of subject areas, we can draw on a rich repertoire of tools and approaches from psychology[50-53], robotics[61-63], and neuroscience[64, 65] for investigation of AP in cell biology.

There is an initiative among leading biologists to reframe the notion of what it means to explain the causes of a biological event [66]. It has been suggested that the classic feed-forward view based on molecular and genetic pathways, particularly in the context of developmental biology, falls short of explaining the adaptability and plasticity of large scale anatomical pattern formation even when they accurately describe the interactions of the individual molecular components [9, 67, 68]. Recent proposals prescribe the basal cognitive lens [69]-a strategy based on exploiting the deep parallels between anatomical pattern regulation and aspects of cognition such as perception, memory, and learning to better model and ultimately control these processes. Thus, this study fits neatly into the paradigm of basal cognition, framing the anatomical pattern formation in angiogenesis in terms of perception, action and behaviour.

## Results

### 1. Filopodia speed up tip cell selection *in silico* and *in vivo*

Early simulation predictions with the memAgent model [24] predicted that filopodia and Vegf gradients alter selection speed. Significant additions and extensions were rapidly made to this primitive agent-based model to incorporate a spring mesh to better model cell shape changes of the cell membrane by the actin cortex beneath [25]. Thereafter, the model was referred to as the memAgent-spring model (MSM) and has been widely used and extended to explore many other roles of filopodia and cell-cell interaction mechanisms during angiogenesis, with a wealth of predictions validated in vivo[26, 27, 70] [71]. Here, we present the first confirmation and extended study showing the MSM model, with its more realistic and verified cell shape changing spring mesh, recapitulates the precursor models first predictions. In order to answer the question “do filopodia impact Notch selection timing” we constrain the MSM model to simulate only the very first, small rapid protrusive filopodia movements of ECs that are otherwise fixed in space, mimicking a first pre-existing vessel stimulated by Vegf in a tissue (see Methods -MSM model).

We found that even small changes to the parameter determining the probability of extending a filopodium (F) in the model (initiating and new one or continuing to extend an existing one) resulted in a delayed selection of the tip /stalk alternating “salt and pepper” pattern (Fig. 2a). F was normally set to 2 in all published works with this model. To assess loss-of-function (LOF) and gain-of-function (GOF) effects, we varied this value in increments of 0.5 in the range [0:3] with the most distinct changes to selection timing found with the values 1 and 3 (Fig 2b, c). To be clear, by LOF and GOF we mean partial rather than total reductions and increases in function. We define the “time to pattern” as the time until certain criteria capturing the salt and pepper pattern are stably met (see Methods -MSM model). In agreement with the precursor model, a reduction in VEGF gradient steepness or concentration also slow patterning (Fig. 2c and Supp. Fig 1). New simulations predict that GOF filopodia extension can rescue patterning speed in LOF VEGF conditions (Figs. 2c, 2d). This points to an intrinsic interrelation of filopodia effects on tip selection speed with the quality of local activating signal in their environment.

**Figure 2.**
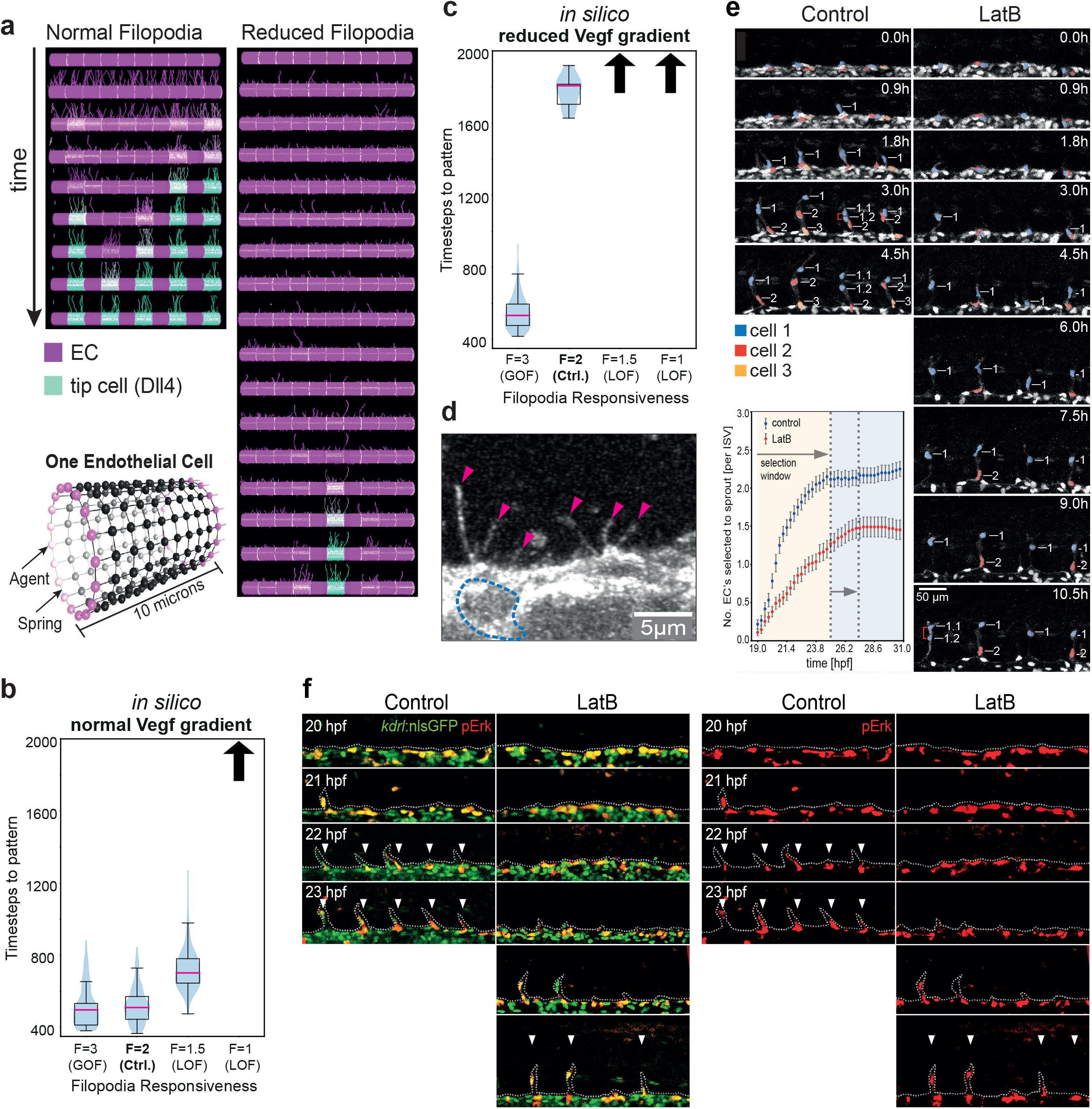
Filopodia LOF slows down Notch Selection: **a)**Time-space plots showing simulations of the MSM model with ten cells fixed in place in a pre-existing vessel, as they detect vegf, extend / retract filopodia and select an alternating ‘salat and pepper” pattern of tip /stalk cells via VEGF /Notch lateral inhibition. Frames shown every 100 time-steps vertically (Dll4 level high-low shown green-purple). Halving the probability of filopodia extension (F = 1 ‘reduced filopodia’) slows down the collective patterning. In the simulations shown, only one tip is selected in twice the available time window compared to 5 with normal filopodial activity. **b**,**c)** Violin plots showing that increasing filopodial activity (gain-of-function, GOF) speeds up patterning in silico both for normal and reduced VEGF gradient and decreasing filopodial activity (loss-of-function LOF) slows down patterning. Blue shading corresponds to estimated probability density of timing values in that range. Upward arrows indicate pattern not achieved within the time window of the simulation for the corresponding parameter settings. Pink line indicates median value, black lines indicate middle quartiles, whiskers indicate upper and lower quartiles. N=100 simulations ran for 2000 timesteps. **d)**A representative image of a cell in the dorsal aorta (DA), showing filopodia formation before tip-cells have been selected to leave the DA into the ISV. The dotted blue line highlights the nucleus of the EC, pink arrows mark filopodia. **e)** Time-lapse images of sprouting ISVs in control and LatB-treated Tg(kdrl:nlsEGFP)zf109 embryos from 19 hpf, and quantification of the number of ECs selected to sprout per ISV over time. Red brackets indicate dividing cells. Nuclei are pseudocolored. Fewer tip cells are selected to sprout in LatB-treated embryos and the selection process is slower. Scale bar, 25 µm. n = 2 independent experiments (55 ISV tip cells were quantified from 15 embryos). **f)** Lateral views of pErk immunostaining (red) in ECs (green) of WT LatB-treated Tg(kdrl:nlsEGFP)zf109 embryos from at the indicated stages. Arrowheads indicate patterned pErk-positive tip cells. Tip cell patterning is delayed in LatB-treated embryos. N=2 (one experiment shown).

To provide the first validation of these prediction *in vivo* we analysed tip selection in the dorsal aorta (DA) of zebrafish embryos. We chose developing zebrafish in particular because they are 1) easily amenable and well-validated for live imaging angiogenesis being transparent during development[72] and 2) sprouting of ISVs from the DA is a highly stereotypical process with simple trackable branch initiations arising from a single pre-existing DA vessel[47], very similar to the simulation setup used above.

First, using high spatiotemporal resolution imaging of emerging tip cells in live zebrafish embryos, we confirmed our time-ordered argument to be correct, as filopodia were observed on ECs residing in the DA at timepoints prior to tip cell selection and ISV branching (Fig. 2d). We then used low-dose latrunculin B (LatB) from these early timepoints to specifically disrupt filopodia extension by perturbing F-actin nucleation at filopodia (but not other actin nucleation sites) [27, 44]. This severely perturbed the speed of tip-stalk decision-making. Unusually, it also extended the otherwise stereotypical time window for tip cell selection (albeit by only 3 hrs; Fig. 2E). The extended time window was akin to observations in Dll4 deficient embryos, suggesting that the selection window is determined largely by filopodia and Notch, but not by Vegfr signalling [20]. Moreover, quantification of the number of ECs residing in the DA that are selected to branch revealed that a loss of filopodia severely restricts the number of tip cells selected to enter branching ISVs (Fig. 2e). Hence, these observations were consistent with a key role for filopodia-mediated temporal control of the collective decision-making process.

Next, we wanted to assess whether the reduced rates of tip cell selection observed in the absence of filopodia (Fig. 2e) were associated with defects in lateral inhibition and establishment of the salt and pepper patterning of Vegfr-active tip cells. To observe tip cell patterning, zebrafish embryos were immunostained for pErk, a highly-specific readout for active Vegfr in EC tip cells [20, 27, 73], at the indicated time points during tip cell selection (Fig. 2f). In control embryos, Vegfr-active ECs were abundant and heterogeneously spread throughout the dorsal aorta during early phases of tip cell selection (up to 21 hpf). This broad and random distribution then rapidly resolved to a refined, highly regular salt and pepper pattern of Vegfr-active ECs by 22 hpf, indicating robust tip cell patterning coincident with initiation of ISV branching, as previously described [74]. Consequently, tip cells that initiated ISV branching were consistently pErk-positive (Fig. 2F).

In contrast, the tip cell patterning process was delayed in latB-treated embryos lacking filopodia (Fig. 2f). In particular, the earliest patterning step whereby ECs exhibited heterogeneous Vegfr activity throughout the DA was prolonged by up to 3 h in latB-treated embryos, with non-patterned clusters of pErk-positive cells evident until 24 hpf and longer. Importantly, this perturbation of tip cell patterning was not due to an overall embryonic delay (data not shown).

Hence, *in vivo* observations were consistent with simulation predictions that filopodia speed up the collective decision-making process. Likewise, tip cells were slower to emerge from the DA and lead ISV branches (Fig. 2f), in keeping with the time ordering of events predicted in the simulations, whereby (1) Vegfr is initially activated in many cells in response to Vegf before (2) slower commitment to tip identity and behaviour defined by filopodia-regulated Dll4-Notch lateral inhibition. Hence, filopodia may influence the speed of tip cell selection to ensure blood vessel branching occurs within the precise time constraints of the developing tissue.

### 2. Filopodia generate bistable switches in individual EC decisions

The ability to dynamically select between two distinct phenotypes while intermediate states are unstable is a hallmark of bistable regulatory systems. In regulatory circuits, bistability is a mechanism well documented to enable continuous systems to act like discrete switches, by destabilizing possible intermediate values of the relevant state parameters in all but the two stable configurations[75-78]. Bistable systems are robust because small perturbations around one of their steady states simply lead to the system returning to its original configuration due to the inherent dynamics involved.

In a vessel comprised of ECs with bistable tip /stalk properties, the attractiveness of choosing one state or the other would enable them to come to a collective patterning decision faster by avoiding drawn out battles in intermediary states. We recently identified that positive feedback to the Vegf sensory system by TM4SF18 slowed selection with indications that this was due to bistable properties [20]. Hence, we seek here to evaluate the hypothesis that positive feedback specifically caused by filopodia movements up a gradient of Vegf speed up selection by conferring bistable decision dynamics to individual cells using a specifically adapted MSM in silico modelling setup, building on preliminary studies [79].

To explain our approach and expected results when assessing this hypothesis, we will use a simple mechanical analogy of a bistable system. The analogy is of a metallic ball found on a double-well landscape (Fig. 3a). In this case, the physical shape of the landscape supporting the ball against gravity creates two stable (attractor) states, i.e., the bottom of each well. In the presence of some small noise able to overcome locally stable bumps in the surface, this mechanical system will not linger in an intermediate state at the top of the barrier or on its slope. Transitions between the two locally stable states take place abruptly, whenever forces on the ball exceed the potential barrier imposed by the presence of gravity. Moreover, slowly tuning the environment by moving a magnet positioned underneath the track left to right and back again (such that the ball is restricted to the left / right / left valleys) uncovers a signature behaviour of bistable systems, namely hysteresis (Fig. 3b). Charting the position of the ball as a function of the position of the magnet clearly reveals the initial-condition dependence of the ball’s response. For any position of the magnet within a window around the barrier, the ball can settle into either of the two valleys, but which one depends on its recent history of positions (Fig. 3c). It cannot be stable in the transition state on top of the barrier, it will fall into one of the valleys. In a system without hysteresis, such as for example if the landscape was completely flat (Fig.3b), the balls resting position would be independent of its past values (depending only on the instantaneous value of the magnets position) and two curves would be perfectly overlapping each other (Figs 3d). Our hypothesis, framed this way, would implicate filopodia in the generation of the hill in the middle of the hypothetical “landscape” in this analogy (Figs. 3e,f,g).

**Figure 3.**
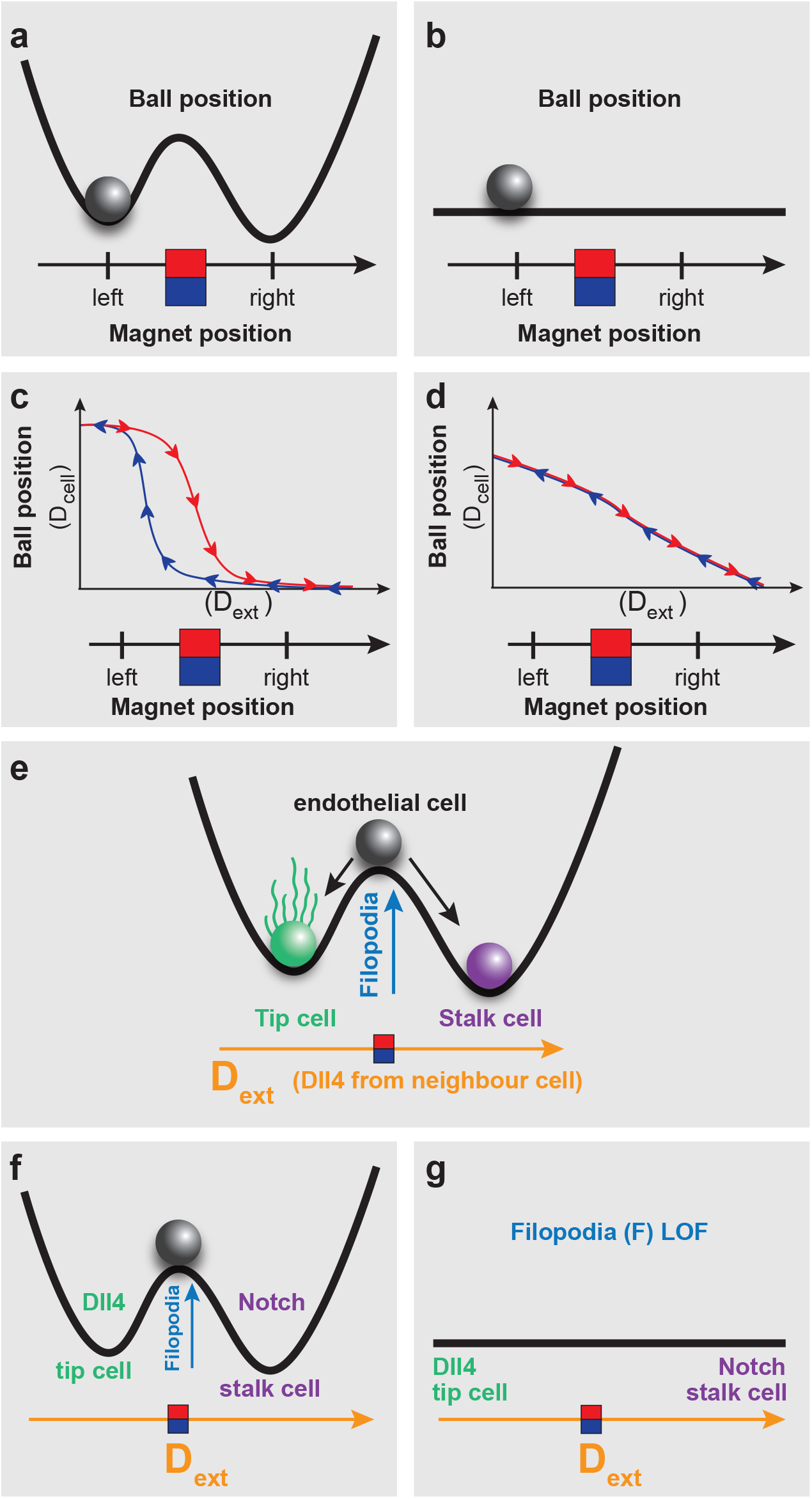
Hysteresis and Bistability. **a)** Metal ball on a double-welled landscape driven by a magnet: analogy for a bistable system, see text for explanation. **b)** Metal Ball on a flat landscape driven by a magnet: analogy for system that has lost bistability, see text for explanation. **c)**Example bistable plot with hysteresis, d) shows the response variable Ball Position (BP) as a function of the driving variable Magnet Position (MP). Of note is the “gap” between the dynamics of BP as MP is moved left (blue line) and the dynamics of BP as MP is moved right (red line) - a typical characteristic of systems with hysteresis. **d)**BP dynamics as a function of MP in an example scenario where there is no bistability or hysteresis. Note how the state variable BP linearly responds to the driving variable MP. **e)**When this analogy is applied to Notch selection in EC’s, the two stable valleys in the landscape correspond to the Tip and Stalk states, and the barrier between them is hypothesized to be caused by filopodial activity. The external magnetic driving force is analogous to the external Dll4 received by the cell from its neighbours. **f)** In the case of normal filopodia, the landscape is hypothesized to have a double-well. In this case we expect the dynamics to resemble that of c. **g)**In the case of reduced filopodial activity (LOF), the landscape is hypothesized to flatten. In this case, we expect the dynamics to resemble that of d.

To assess whether filopodia confer the bistable property of hysteresis to individual EC decisions during selection, we established a modified version of the MSM tip cell selection model designed to perform hysteresis analysis on a single EC (see methods-MSM-hyst). One crucial difference to the model here is that though we simulate three cells, we only test the dynamical properties of the single central cell. The two “cells” either side of the central EC in the model don’t act or respond as ECs, but merely provide fixed sources of external Dll4 that the central cell can bind (*D*_*ext*_) akin to the magnet in the mechanical analogy. (Fig.4a). When the side cells have a fixed high level of Dll4, the central cell becomes inhibited via this Notch-Dll4 signalling prohibiting it from becoming a tip cell (Fig. 4b). Conversely when the side cells are fixed to present low Dll4, the middle cell receives no Notch signal and extends filopodia, becoming a selected tip (Fig.4c). To assess hysteresis in simulations, *D*_*ext*_ starts at 0 and is then increased in small increments up to *D* _*max*_ then decremented back down to zero, akin to the magnet movement left and right.

**Figure 4.**
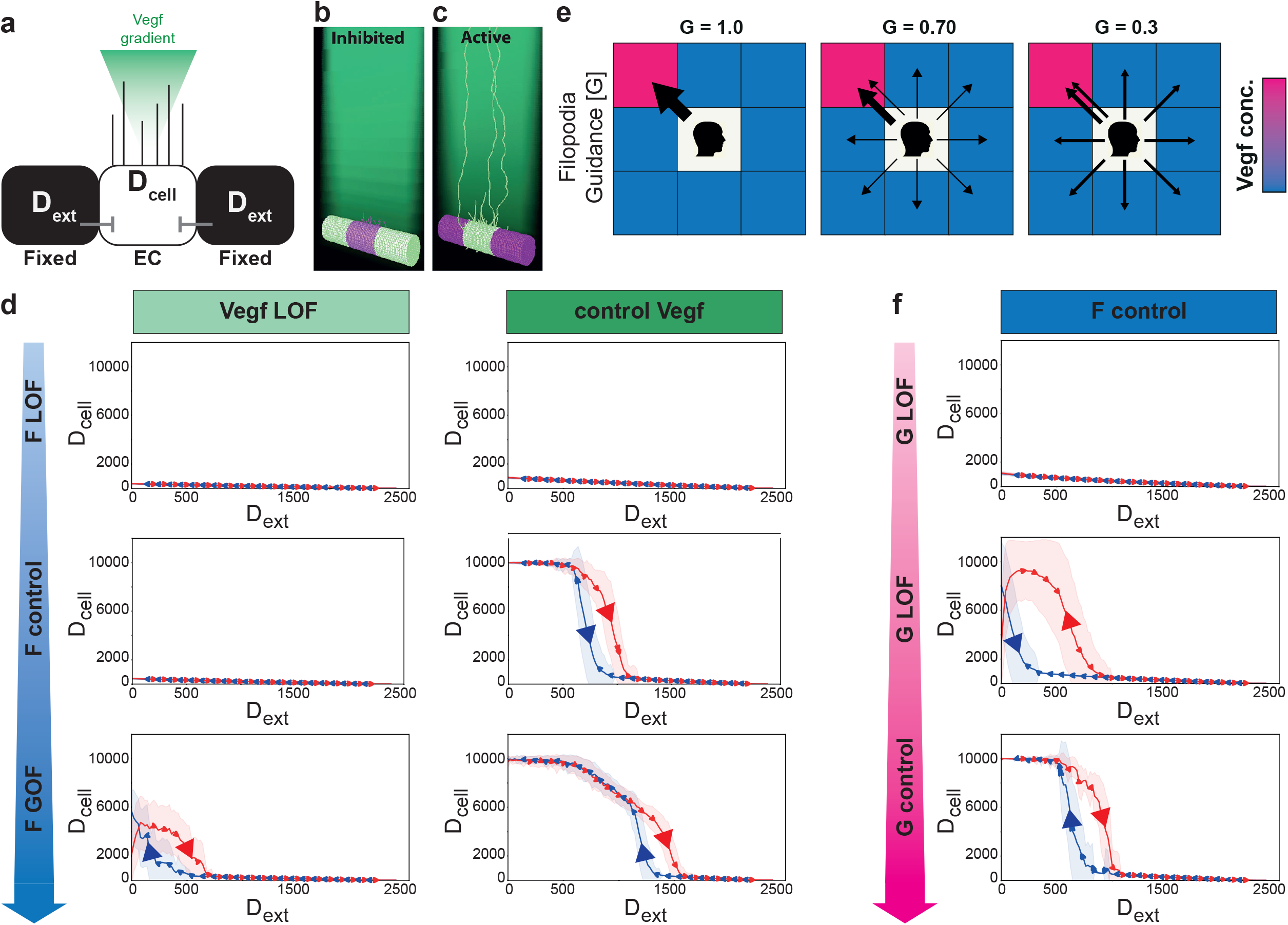
Filopodia Movement in Vegf gradients confers bistable properties to Notch patterning. **a)** Schematic of the MemAgent-Spring Model Hysteresis (MSM-Hyst) testbed, designed to probe the behaviour of the single endothelial cell in the middle (through *D*_*cell*_ dynamics) in response to externally driven inhibitory *D*_*ext*_ signals from two neighboring pseudo-cells and VEGF in the environment. **b**,**c)** Screenshots of the MSM APB simulation testbed when *D*_*ext*_ is at max (b) and min (c) levels. b) the central cell is inhibited, c the central cell is uninhibited so has high Dll4 and filopodia (tip cell selected). Key: Dll4 high cells - light green, Dll4 low cells – purple, VEGF gradient in the environment - dark green. note the peripheral pseudo-cells do not have filopodia, as they only provide *D*_*ext*_, they do not to respond to vegf or Notch themselves, in this highly constrained testbed. **d)***D*_*cell*_dynamics as a function of *D*_*ext*_in cases where we covary the model parameters determining the steepness of the Vegf gradient (0.02(LOF), 0.04(control)), and the responsiveness (in terms of probability to extend) of the filopodia to sensed Vegf(1(LOF), 2(control), 3(GOF)). We can see that some parameter settings (e.g. F=2, Vegf gradient = 0.04) yield bistable dynamics with hysteresis, while others (e.g. F=2, Vegf gradient = 0.02) do not. N=100. Shading indicates 1 standard deviation. **e)** A cartoon demonstrating how the filopodia guidance parameter (G) would influence the behaviour of a growing filopodium in a 2D schematic (the MSM simulation extends this to 3D). The central square represents the memAgent at the head of the filopodium, who must decide which direction to step into every timestep that the filopodium extends (see methods for full details). G determines the probability that the memAgent will choose the adjacent square with the highest level of Vegf - Vegf high (pink), Vegf low (blue). Thickness of the arrows represents the probability of stepping into a given square. When G=1 (left panel), there is a 100% probability of stepping into the pinkest square. As G is decreased, the probability of stepping into a square at random increases - represented by uniformly distributed arrows and a thinner arrow pointing to the pinkest square. Note that the probability of stepping into the pinkest square is never zero, as stepping at random might still land on it. **f)** *D*_*cell*_dynamics as a function of *D*_*ext*_ where we vary the guidance parameter G. As G approaches 0, the hysteresis window shrinks and the bistable dynamics disappear. For all plots, N=100, shading indicates 1 standard deviation. F = 2 and V = 0.04 (both control settings).

To evaluate the hypothesis that filopodia moving through a Vegf gradient generates positive feedback in the Vegfr system and confers bistability, we disaggregated that idea into parts and performed simulations varying each part. First, we covaried the probability of filopodia extension (F) with the slope of the linear Vegf gradient (*V*) and analysed the Dll4 dynamics of the middle cell (*D*_*cell*_) as *D*_*ext*_ levels were incremented up and down. In the ‘normal Vegf gradient’ (V = 0.04) with standard Filopodia responsiveness (F=2) both calibrated previously and used throughout MSM studies [24]) a bistable window is indeed apparent (fig 4d, control Vegf).

When F is halved under normal environmental conditions (F=1, V=0.04), the reduction in filopodia movement eliminates the bistable properties of the MSM-Hyst cell (Fig. 4d). Similarly, when the steepness of the Vegf gradient is halved (V=0.02, LOF), the bistability is lost (F=1,2) - though the hysteresis is partially rescued by GOF in filopodia movement (F=3)(Fig. 4d). In cases with high filopodia activity and positive feedback from a steep Vegf gradient, such as *F* = 3, *V* = 0.04, the curves coalesce near the top of the bistable region, but we believe this to be a negligible artefact of the boundary conditions we use in the model (if the filopodia manage to reach the gradient ceiling, they have nothing left to climb and lose their positive feedback, thus disrupting the hysteresis and bistability and reverting to linear response behaviour).

As a reduction in Vegf gradient steepness incurs a knock-on reduction to general Vegf concentration in this model (see methods – MSM model), this could be causing the very low *D*_*cell*_ values and a loss of bistability simply due to insufficient Vegf to initially activate the cells in the Vegf gradient LOF condition (Fig. 3d). To control for this, we utilised a previously calibrated version of the model with no gradient (a uniform distribution of Vegf) but using a concentration high enough to stimulate cells to achieve a pattern. Using our precursor memAgent model, we previously reported a slower time to select when Vegf is distributed uniformly compared to in a gradient [24] and indeed we see bistability is lost in these conditions in the MSM model regardless of the level of F and the concentration of VEGF in the environment (supp. Fig.2).

Taken together these simulations predict that positive feedback to VEGF sensing by means of filopodia movements through a Vegf gradient do confer rapid and robust bistable switch-like properties to single ECs experiencing changing dll4 conditions from their neighbour cells.

Next, to test the importance of guidance up the Vegf gradient, as opposed to merely Vegf levels and filopodia responsiveness, we uncoupled the intrinsic feedback in the MSM model between Vegfr (sensing) and filopodia extension (movement). We did so by modulating the guidance parameter *G*, which denotes the probability that a moving filopodium will choose to extend in the direction up the gradient - else taking a random step (Fig.4e). *G* ranges from 0 → 1, with its default value set to 0.9 based on experimental observations [43].

By varying the Guidance parameter (G) we found bistability properties are visibly disrupted with lower values, i.e. when the movement is less coupled to sensing (Fig 4f, *G*= 0.3) at which point the active state loses stability. Though this low threshold value appears to minimize the importance of guidance in the hysteresis properties, the simplicity of the linear vertical gradient in the square lattice-based model means that a filopodium would step in the correct direction of positive feedback approximately 1 /3 of the time even if it is moving completely at random. When adjusted for this fact, we see that even a 40% reduction in effective guidance (from 93% adjusted to 53% adjusted) is sufficient to diminish bistability. This is comparable to the changes in F-where cutting the value in half disrupted bistability and hysteresis. Conversely, GOF had no significant effects but given that G is normally close to its maximum value this is to be expected.

In summary, altering parameters involving the guidance (G), filopodial responsiveness (F), or the gradient itself (V) can cause an abrupt transition to non-bistable tip /stalk dynamics. The bistability in this system is contingent on all three components being present - filopodia movement, the presence of a gradient, and the ability of the filopodia to navigate up the gradient.

### 3. Measures of ‘enriched’ sensation by coordinated motion suggest filopodia are a basal form of AP

Our results so far indicate 1) a role for filopodia in speeding up collective tip cell selection *in vivo* (fig. 2) and 2) that positive feedback between sensing and movement via filopodia confer bistable switch-like properties to facilitate rapid decision-making (figs. 3,4). Here we ask the question are filopodia, therefore, a form of active perception? Are they akin to a basal cognitive function exploiting sensorimotor coordination to enable endothelial cells to adapt their function to the changing local environmental conditions by making ‘better’(faster, more efficient) decisions?

To answer this question, we adopt the formalism of Lungarella, Sporns, and Pegors who define active perception as the enrichment of the statistical structure of an individual’s sensory input resulting from coordinated sensory and motor activity [64, 80, 81]. To detect active perception, they suggest a series of measures to probe statistical structure in a compressed representation of sensory input (the ‘Sensory Map’) induced by coordinated sensorimotor activity. They found that the measures they used have distinctive signatures, such as reduced Shannon entropy when their model systems are allowed to coordinate sensory and motor activity [64].

In order to apply this approach to ECs, we first established a simple two-cell version of the *in silico* MSM model, as this is the minimal number of cells required to achieve the patterning outcome, and thus the simplest first case to delineate AP in a collective decision (Fig.5a). We here identify the cells reaching a “pattern” by comparing their Dll4 levels in order to pinpoint the earliest timepoint that Notch selection has stably occurred (in contrast to the longer-term Vegfr and filopodia dependent “time to pattern “score used previously). A pattern is recorded when one of the two cells has obtained and maintained a substantial lead in Dll4 for a long enough time (see methods -MSM hyst.). The cells were again fixed in space and only able to extend and retract filopodia into a VEGF gradient, as per all simulations in this study.

We define our Sensory Map to be the projection of all activations of the Vegfr-2 receptors of both cells onto the fixed cylindrical vessel surface (Fig.5b). Receptor activations occurring on a filopodium are projected from their point in 3d space down to its nearest point on the cylindrical shell; this mapping preserves directional information (as opposed to collapsing activations down to the filopodium’s base position on the cell surface) and results in a fixed spatial representation of the sensory input that the cells experience, overcoming the challenge of measurements on a dynamically changing form. The sensory map is considered at the scale of the vessel rather than the individual cell, i.e. we are specifically asking the question here does the vessel employ AP to solve a collective cell decision problem? Though this representation necessarily loses some information, we hypothesized the sensory map would retain enough directional information to detect a difference in the measures. This approach considers the question of AP at the scale of the vessel rather than the individual cell.

To search for signatures of active perception, we calculate the following measures across the sensory map at each time step in the simulation:

1. **Intensity (I)**: the sum of Vegfr activations (Vegfr*) across the sensory map with dimension N. This value is in units of Vegfr activations.

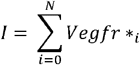
2. **Shannon Information Entropy (H):** of the spatial distribution of Vegfr activations, for which we infer a distribution by binning with Laplace smoothing [82]. This is in units of Bits (basic unit of information, here quantifying uncertainty).

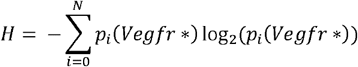

Where *P*_*i*_ is the inferred probability density of the amount of Vegfr activations at the i’th site of the sensory map.

Rather than extensively testing for any possible structure in the sensory map, we chose these AP signature measures in particular because changes in these would be straightforward to interpret yet form a basis and approach easily extended to other measures in the future. Shannon Entropy (H), has a particularly useful property – it increases for broader distributions. E.g. a sharply peaked Gaussian distribution will have a lower entropy than a uniform distribution over the same space. Thus, we can use the value of H to describe the spatial spread of activating receptors on the cell surface (Fig. 5c). Crucially, this corresponds to the goal of the tissue as a whole to adopt an inhomogeneous pattern - which makes H a quantitative readout of progress towards this goal.

**Figure 5.**
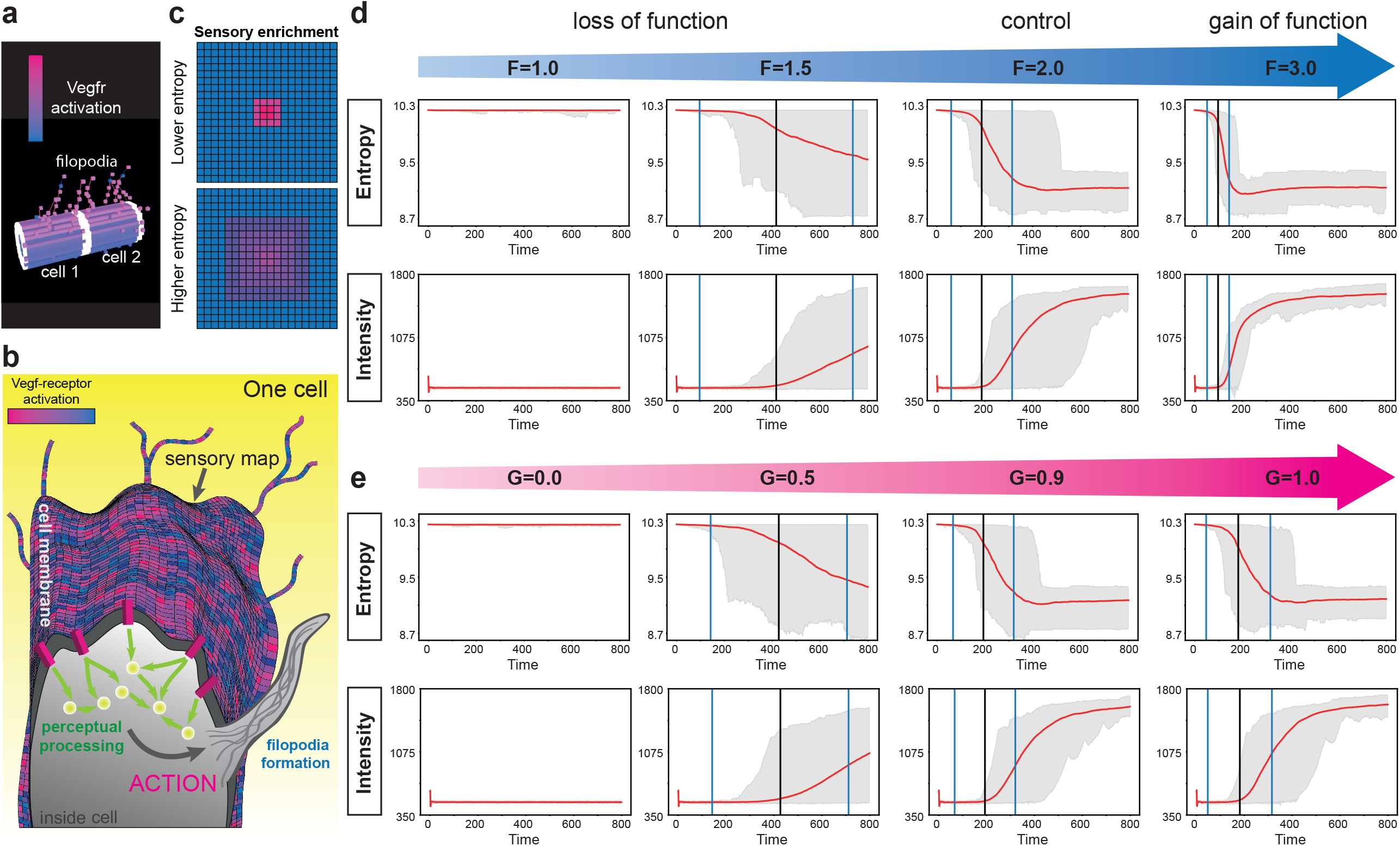
Signatures of Active Perception. **a)** Two-Cell MSM simulation setup used for active perception signature studies. Sections of cell membrane (MemAgents) with higher VEGFR Vegfr activation– pink cubes, memAgents with low Vegfr activation - blue cubes. Structures extending out are filopodia. White cubes are memAgents on a junction with the other cell (indicating periodic boundary conditions far left and right). **b)** Cartoon diagram of a sensory map. A sensory map, in this study, is a coarse-grained representation of what is sensed by endothelial cells. Activations of the Vegf receptors on memAgents throughout 3D space (e.g. on filopodia) are mapped down onto the cell surface. All AP measure analysis is performed on this fixed spatial representation of Vegfr activations changing over time. **c)** Example sensory maps with high and low entropy Vegfr activations (pink – high activation). Because the sensory map itself is surface of a cylinder (due to the shape of the cell surface in simulations), we may imagine it “unrolled” as a 2D rectangle, having various Vegfr activations across it. In the sensory map labelled “Lower Entropy”, the Vegfr activations are highly spatially concentrated in comparison to the “Higher Entropy” case where they are more diffuse - yet their total intensity is the same. **d)** Active perception measures as a function of time for different values of F, the filopodial responsivity parameter. Red lines are the mean values of N=100 runs of the 2 cell MSM model. Grey shaded area is the entire span of the data set. Vertical black line is the mean time at which the salt and pepper pattern was achieved (counting only the cases where a pattern was achieved within the max timesteps, up to N=100), vertical blue lines bound the regions within two standard deviations of that mean. **e)** Plots showing time evolution of measures for different values of G, the parameter determining filopodial guidance.

We hypothesized that signatures of active perception would be apparent in the time evolution of the measures applied to the sensory map. For example, a reduction in the entropy of the sensory map over time would mean the driving Vegf signal that the vessel is receiving is being biased to be less spatially homogeneous across it. In the beginning, both cells have an equal chance of “winning” and becoming the tip cell (this is due to stochasticity in the simulated filopodia extension process). Ultimately, the cell that wins is determined by the sensory input that the pair receive. If one cell starts to get more of the growth signal, this would both facilitate the production of an inhomogeneous pattern (e.g. more sensory input localized on one region of the vessel – the winning cell) and correspondingly reduce the measured entropy of the systems sensory map. If active perception is driving this entropy decrease, then the disruption of active perception would reduce or remove the reduction in entropy.

In terms of the intensity of Vegf activations across the vessel, we expected that the intensity as a function of time takes on a different form, for example by switching from a linear to a nonlinear regime due to increased feedback between movement and sensing. Overall, we expect a filopodia (LOF motor response) in a low Vegf gradient (LOF stimulus) would have a reduced AP ability to one with a stronger coupling between sensing the Vegf (V) and the motor response to extend filopodia (F) (i.e. F and V normal or GOF).

As before, we varied the probability of extending filopodia (via F) and the steepness of the Vegf gradient / concentration (V) to see how these measures change. There are marked differences between the sensing and motor response LOF and GOF cases, as shown by the behaviour of the measures as a function of time (Fig. 5d). In the normal and GOF of filopodia conditions, Entropy appears to decline slightly before a stable pattern is achieved – and then goes down by approximately one bit after the patterning is achieved. The fact that entropy decreases before the patterning is detectable in Dll4 levels, however, may be susceptible to the choice of pattern Dll4 criterion parameter (see methods - AP Signatures). Intensity takes a sigmoidal shape as a function of time (Fig. 5d). In the cases where filopodia (actions) are inhibited (F LOF), entropy declines less sharply or not at all and the sigmoidal shape of the Intensity starts to flatten and ultimately disappears (Fig.5d). This implicates filopodia in the generation of these patterns in the sensory map.

A reduction in entropy indicates that the amplification of differences in the sensed Vegf signal have begun. A reduction in spatial homogeneity of the driving sensory signals is evident *before* the patterning decision at the level of Notch signalling is completed (Fig. 5d), thus sensory map structuring is predicted to bias one cell to win before lateral inhibition commits this choice. A sharper reduction in entropy corresponds to a faster patterning decision.

Sigmoidal shapes in the Intensity plot (Fig.5d), on the other hand, commonly result from variables undergoing positive feedback, where the rate of change of some value is positively proportional to its current magnitude. Hence, the shape of the time evolution of the sensory map may indicate the positive feedback facilitated by filopodia. In the previous section, simulations predict that positive feedback from filopodia generates hysteresis and bistability in EC states. Here, by analysing through the cognitive lens we have found evidence of this positive feedback in the time evolution of the intensity measure of Vegfr activations.

When we alter the guidance parameter (*G*), we see that the AP signature changes in line with F or V LOF (Fig.5e). In the normal and GOF cases (*G* = 0.9,1.0) the same pre-emptive, slight reduction in entropy before the Notch patterning decision is made, followed by a drop of 1 bit after. Intensity also responds to GOF in the way one would expect, with a clear sigmoid in the case of *G*= 1.0 (GOF), and a less clear sigmoid for *G*= 0.9. In the LOF cases where *G*=0.5,0.1 we notice the partial, and complete loss of these observed structures. This suggests that the signatures we measured correspond to the effects of active perception as a whole, rather than just one of its components.

Taken together, we find entropy and intensity of Vegfr activations on the sensory map can be used as a signature to connect the effects of the three key component parts of the system under the unified, basal cognitive conceptual umbrella of “basal Active Perception”. When Active Perception is present, the entropy of activations on the sensory map is reduced before a patterning decision is observable in Dll4 levels, and the intensity of activations over time has a sigmoidal form. These findings correspond to ways in which the cells direct their receptors to facilitate rapid and robust patterning.

## Conclusions and Discussion

Here we present the first proof of concept *in silico* and *in vivo* study supporting the theory that endothelial cells utilize filopodia as a form of sensorimotor feedback (Active perception) to speed up Notch decisions during angiogenesis. This highlights the functional importance of the movement of Vegfr through the environment via a deforming surface, rather than merely its basic functionality as a receptor. We find *in silico* that the shape-changing movements of filopodia provide positive feedback to Vegfr-Vegf signalling, which endows the tip-stalk selection machinery with bistable, rapid and robust switch-like properties and hysteresis. Lastly, we begin to unpack the notion that filopodia on ECs may constitute a basal form of active perception. We apply information theory measures used in computational neuroscience, where AP is more well-studied, which indeed indicate that this system meets a necessary but not sufficient criterion for filopodia “enriching” sensory input is met, providing the first rigorous step towards understanding and analysing the way sensor-motor feedback contributes to EC adaptive behaviour during angiogenesis.

Our simulations identify three critical components to the system that ensure ECs are endowed with bistable properties and complete selection rapidly – filopodia, a Vegf gradient and a direct coupling mechanism between them (e.g. Guidance). However, our simulations predict that a gain in one of these components can rescue bistability and selection timing given a reduction of another. This strongly suggests the importance of the interlinked nature of these components rather properties of the individual parts in isolation. In tissues with very shallow gradients or with minimal guidance coupling – e.g. more randomly directed filopodia as in zebrafish ISVs [44], a GOF adaptation of filopodia extension could permit ECs to nevertheless keep selection occurring on time within tissue time constraints.

Our work raises an important new role for filopodia in ensuring the Notch selection of tip cells occurs in a timely, rapid and robust manner. Contrary to the prevailing literature that considers the role for filopodia being *after* tip cells are selected, here we show that they play a key role *before*: Movement must be primary and part of the decision process rather than secondary, after the decision has been made. We thus propose a shift in perspective to considering even small cell shape changing movements as potentially critical parts of the cellular repertoire of tools, helping them to solve complex problems during tissue morphogenesis.

The conventional way to explain biological events is at the scale of molecules, proteins, and genetic pathways. But in tandem to this well-trodden explanatory framework we find viewing events and processes in terms of active perception, through the perspective of basal cognition, brings great benefits for revealing hidden mechanisms in cells for two main reasons:

1. Investigating AP naturally brings embodiment (the cells body shape and organisational material structure) to the forefront. Often, we can forget the spatial shape of cells when reducing mechanisms to molecular /genetic pathways[83]. We recently found that tip cell ECs manipulate their body shape and size during sprouting in order to bypass slow Notch selection altogether and rapidly assign one daughter (the larger cell) to be the tip [27]. This work shows that cell embodiment factors (such as shape, size and movement) can have the same importance as molecular and genetic factors in a developmental mechanism. Actively emphasizing how these dynamically couple to molecular pathways opens the door to a wealth of unexpected insights and hidden knowledge that ‘well-studied processes’ considered only from a non-spatial, molecular /genetic perspective may have missed. The field of biomechanics, putting force changes on an equal footing with molecular signalling by viewing cells simultaneously from a mechanical and molecular perspective revealed a multitude of hidden abilities of cells [84].Yet, we are only just scratching the surface of how the dynamics of cell shape and geometry feedback to signalling[85-87].
2. Pathways are a type of model for understanding biological events at a specific molecular spatial and temporal scale. Despite the tremendous successes of pathways centric biology, it is necessary to build on them and ask broader questions involving feedback, control, and information flow if the goal is to understand and control biological outcomes at the higher scales, such as organism /tissue-level patterning and complex behaviour [67, 68]. For example, if one endeavours to teach a rat a circus trick, it is advantageous to train the rat using positive and negative reinforcement rather than trying to micromanage every individual neuron in the rat’s body [67]. Controlling morphogenic outcomes that result from the collective patterning decisions of cells may similarly benefit from such a behaviour-level approach, which the emerging basal cognition field can provide.

An immediate benefit of framing the tip selection process in terms of AP is the number of new and exciting questions and research directions that open up. Below we briefly highlight a selection:

### Open questions arising from the basal AP perspective

#### Do other cell shape-changing movements form part of basal AP mechanisms?

Here we focussed on filopodia as their presence in angiogenesis in widely characterised, yet their role has remained unclear. However, ECs also exhibit lamella-type extensions and triangular veils [45] [25]. These protrusions are known to play roles in cell migration but a role in enriching sensory input to alter timing of cell decision-making (basal AP) has not been considered. Both work at a marginally slower timescale to filopodia and are generated by cross connected actin rather than parallel bundles. Furthermore, the shape change is fundamentally different, these are more convex or triangular respectively than the long thin filopodia. Altogether meaning that shape-signalling feedback could vary greatly on each kind of cell protrusion, potentially providing a variety of basal AP mechanisms for cells to combine when enriching sensory input for problem-solving.

#### Do ECs continue to use basal AP while migrating and sprouting?

Here we focussed on the very first, small rapid protrusive movements of ECs that are otherwise positionally fixed in a pre-existing vessel detecting a Vegf signal, before the longer-timescale full body movements of tip cell migration during sprouting or EC tip and stalk position-changes within vessels (‘cell rearrangement’) have begun [34, 35]. This was to avoid interference of other roles filopodia or longer time-scale full-body movements in this first study. However, Notch signalling continues after the first tip cells are selected, and the maintenance of a differential salt and pepper Notch pattern is required to generate differential adhesion along the sprout, which drives the cell rearrangements [23]. Thus, the ability to continually and rapidly re-establish the Notch salt and pepper pattern when individual cells change positions must still be required [9, 33]. It is reasonable to hypothesize that filopodia continue to play a decision-making role on migrating ECs within sprouts when the Dll4 signal from neighbours becomes less clear after a local position-switching event disrupts the cell-by cell Notch pattern – akin to the mid-range D_*ext*_ levels in the hysteresis simulations. Indeed, in an integrated in silico /in vivo study of PFKFB3-driven glycolysis in filopodia formation we found that a reduction in filopodia perturbed cell rearrangement dynamics, indirectly by impacting Notch patterning timing.

#### Do ECs utilise basal AP to solve other tasks than selection during angiogenesis?

The study presented here focusses on the tip cell selection step in angiogenesis, which can be seen as a key collective coordination /decision ‘task’. However, there are many different problems cells solve in order to construct the full vascular network that could be posed as ‘tasks’ cells might efficiently solve using shape-signalling feedback, AP, or other basal cognition mechanisms. For example, filopodia are known to improve tip cell fusion during anastomosis [25, 47], but exactly how remains unclear. ECs in formed vessels often exhibit a Cilia – a sole protrusion into the lumen, which aids the sensing of flow through its movement and thus could be a form of basal AP [88].

#### Is basal AP ubiquitous across cell types and tissues?

The really tantalizing question is how widespread basal AP mechanisms like this one may be, among the myriad of cells in tissues of different organisms? Is it an evolutionarily conserved and common mechanism employed by cells to enhance their adaptive problem-solving or a rarity, emergent only if a set of tissue conditions are met? Life-critical processes such as morphogenesis, wound healing and repair are by their nature time constrained, which we can imagine would then favour evolution of basal cognitive processes that can help to solve them faster. Cell shape changes occur as a matter of course during morphogenesis and are currently, largely considered from a biomechanical stance, however they may also confer basal AP properties by enriching sensory input, justifying wider application of the basal cognitive lens in tissue morphogenesis research. A further cell type within organisms that stands out as a prime candidate for basal AP are macrophages as they have distinctively protrusive, dynamical filopodia and lamella, which appear to play multiple roles aiding the cell to rapidly traverse very different tissue environment ‘terrains’ while finding and catching foreign bodies to phagocytose [89].

#### What are the molecular players comprising basal AP mechanisms?

Here we co-modulated the three core interrelated components of basal AP (sensing, movement and the coupling between them) and targeted filopodia as a first coarse-grained approach. The next step is to delineate the precise molecular pathways coupling sensing and movement within filopodia, such as receptor trafficking mechanisms within filopodia and assessing whether membrane shape-detecting proteins like F-BAR [90] may play a role in coupling signalling to the shape changes. Interestingly Tm4sf18, which we found conferred positive feedback to Vegfr sensing and facilitated switch-like rapid selection *in vivo*, has been previously shown to increase filopodia formation [91], making it a prime target as a basal AP component. In a prior study to delineate the role of Cdc42 (a small GTPase involved in the regulation of actin-based morphogenesis and cell polarity [92-95]) in angiogenesis, we found that ECs lacking Cdc42 in the mouse retina *in vivo* have a competitive disadvantage to assume tip positions compared to ctrl ECs in chimeric angiogenic sprouts [96]. Cdc42-KO-EC at tip positions lacked filopodia and instead protruded only few very short and studded protrusions from their surfaces. This suggests a crucial and cell-autonomous requirement for Cdc42 in tip cell selection. If a wider array of basal AP mechanisms were found across tissue contexts, the natural next step is to check if there is a core set of evolutionarily conserved AP molecular pathways - or if cells make use of very different proteins with the same functionality to couple sensing and motion in different contexts.

### Challenges of AP investigation at the cell scale

There are many opportunities but also challenges, and responsibilities to carefully consider, given the impact that basal cognition focused studies such as this could have for the field of cell and tissue biology within higher organisms. One such problem is to balance and temper the psychological concepts and claims in AP studies made with higher organisms correctly, when working at the cell-level. Similarly, there is a need to consider the empirical tools and viability of features of experimental design for study of AP at the different scales.

To establish conclusive evidence for active perception, Yang et al [22] suggest that it is necessary to compare the actions of the system in question to those of a model of what an idealized active sensor would do given the same goal (for example, a model utilizing Bayesian decision theory [55, 97]). Characterizing active perception in our system this way is challenging because doing so would require assuming that endothelial cells have a known goal in their actions. Even if cells are optimizing towards a goal it is often the case that they are simultaneously solving a number of problems in parallel. This makes it hard on teleological grounds to devise a simple idealized model for comparison (except for potentially viewing the negative feedback of lateral inhibition as a form of teleology [98]). Thus, we took a simpler, indirect approach when assessing the presence and effects of active perception in angiogenesis as utilized by the Lungarella et al[80].

We defined, akin to Lungarella et al, basal AP for cells in tissues as specifically meaning coupled, direct two-way feedback between motion and sensing (i.e. not requiring indirect processing through gene regulation) that enriches sensory input leading to improvement of a collective cell problem-solving process in morphogenesis. This minimal definition avoids unhelpful, overfitting of human-level psychological connotations to cells such as perception, action, goals and intentionality while enabling empirical testing of basal AP in the future. This enables us to now use verifiable molecular approaches in vivo and in vitro to prove the coupling between motion and sensing. Together with signature changes to the sensation enriching measures, this is a way to measure the resulting effect on the cell coordination process in question. Here we focussed on the first simplest sensory enriching measures – the time evolution of receptor activations and entropy. We plan to explore and extend to more complex metrics in the future, such as using transfer entropy to assess the strength of sensorimotor coupling rather than just searching for its signatures [64, 99].

Overall, we hope that this study acts as a first step into a new research programme elucidating the extent and variety of basal AP mechanisms that cells from different tissues may have evolved to enable complex collective cell coordination processes. Importantly, as many seemingly unrelated pathways may constitute component parts of basal AP mechanism, we propose that basal AP is inherently accessible to therapeutic modulation by targeting any of these to control and tune individual to collective cell behaviour in tissues.

## Methods

### *In silico* Methods

#### 1. The MemAgent-Spring Model (MSM)

The MSM model was first developed and described in full by Bentley et al in [24, 25] and has since been used in a wide range of studies to explore VEGF /Notch and tip cell dynamics during angiogenesis with many predictions being validated in vivo from this model[26, 27, 70] [71]. Being the only angiogenesis tip selection model with sophisticated filopodia dynamics, it was the natural choice for extension in studies here. Briefly, in this model cell morphology is represented as a surface comprised of many small agents (membrane agents, “memAgents”) connected by springs following Hooke’s law (which confer tension to the changing cell shape, akin to the actin cortex beneath a cells membrane) (Fig. 2b). The memAgents move in continuous space but are snapped to a gridded lattice to implement discrete local rules of agent interactions (grid sites represent 0.5×0.5×0.5-micron cubes of space). At each time-step an equal number of the cell’s current, total Vegf receptors are distributed to each memAgent over the cell’s current surface shape. Dll4 ligands are only distributed to any memAgent in Moore neighbourhoods of memAgents from other cells (on the interface between cells or “junctions”). The ambient Vegf is modelled as a fixed linear gradient or a uniformly distributed value, where in the linear case each grid site (g) starting from y=0 has *I*_*g*_ level, calculated as *I*_*g*_ = *V*_*slope*_ ·*y* where *y* is the *y* axis coordinate position of *g* and V is the slope of the gradient.

##### Model initialization and parameterization

The model was initialized with 10 cells in a row, one per vessel cross section (e.g. Fig 2B), representing a collection of endothelial cells in a pre-existing vessel competing to sprout into the space above (represented very simply here as just a fixed VEGF gradient extending into the y axis above the horizontal row of cells). All parameters were kept the same as previously published [17, 20, 24, 25] except those being varied to match the experimental conditions here, described below. E.g. a cell is 10 microns wide and spans one cross section of a pre-existing vessel with 6 microns diameter (matching *in vivo* capillary dimensions) Each cell comprises approximately 1200 memAgents initially.

##### Signalling

Simplifying the description of the competitive receptor biology between different Vegfr’s in the original MSM model, as it is not the direct focus of this study, we can state that the number of active Vegf receptors of a given memAgent m is calculated as 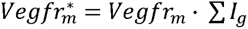 – where the * denotes an activated receptor-ligand complex. Notch is activated in each memAgent (resulting in the local removal of Dll4 from neighbouring cell’s memAgents), up to the value of *N* _*m*_. If Dll4 < *N* then 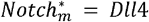; if *Dll*4 > *Notch*, then *Notch*_*_ = *Notch* _*max*_.

##### Filopodia

MemAgents extend filopodia by one grid site length at a time by creating non-branched chains of new memAgents adhered to the environment (fixed in position by adhesions to the environment every 2 microns, as per ([46]). These events are stochastic, with approximate probability:

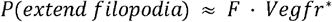

where F is a tunable constant varied in simulations here. Filopodia do not extend unless they have enough available actin and vegf receptors activate. For full rules behind the actin dynamics and spring settings driving the filopodia extension and retraction see [25]. Filopodia retract if the memAgent at the top of the chain has not extended for Fmax. Retraction occurs by iteratively snapping the chain of springs back to the next memAgent in the chain, whilst deleting memAgents as they go (based on *in vitro* observations). The cells have no limit on the length of a single filopodium, or how many they can initiate. However, their total length is limited by the total amount of actin available to the cell, which gets used up on every filopodia extension of one micron. The original model in [25] generates full cell migration by releasing filopodia adhesions and allowing the springs to pull the cell forward akin to veil /lamella advance [46]. Here, however we keep adhesions fixed to control variables, prohibiting these other forms of protrusions, to ask specifically whether shape changes due to filopodia play a role in selection timing.

##### Genetic Regulation

Briefly, Notch lateral inhibition is implemented each time step as follows:

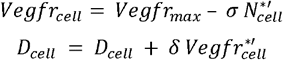

where •, • are constants and *Vegfr*′* denotes Vegfr* after a time delay (28-time steps) corresponding to the time it takes for translocation of activated receptor signalling to the nucleus, transcription to alter gene expression, and subsequently protein levels at the cell surface. The new *Vegfr*_*cell*_and *D*_*cell*_levels are then redistributed to the memAgents and the process repeats on the next timestep.

##### Parameterization

All parameters in the model are set as in Bentley et al (2009), which were previously calibrated to experimental data. To investigate active perception, we focus here on simulations varying filopodia extension (F) and the input signal gradient (V). For normal EC, F = 2 and V = 0.04, Guidance G=0.9

We switch off veil advancing, lamella mechanisms in simulations, create permanent adhesions at the base of filopodia prohibiting cell body movements and switch off cell rearrangement position changing junction-based mechanisms [23]) fixing cells to their initial positions in the pre-existing vessel.

##### Time to Pattern

To assess patterning times (when a clear and complete alternating salt and pepper pattern of tip and stalk cells have been selected) the model was run 100 times for a maximum number of timesteps (2000 for Vegf gradients and 5000 for uniform Vegf). For the 10-cell version of the MSM the pattern must meet criteria as per Bentley et al [24]. Tip cells are defined as cells that have sprouted filopodia and have not been inhibited (as per *in vivo* tip cell characterisation). The patterning criterion is that four or five, non-adjacent tip cells must be active at a given time. The stability criterion dictates that the pattern of tip cells must persist for 100 timesteps (chosen as it allows for a cycle through the Notch pathway to occur which might alter the pattern if unstable). Once these criteria are met, the collective of cells is deemed “patterned” and the time is recorded.

#### 2. The MSM Hysteresis model

To test our hypothesis that filopodia speed up patterning due to the positive feedback of sensing and acting generating bistable properties in single cells, we developed a new simulation testbed extension to the MSM model that parallels the mechanical ball and magnet example of a bistable system (Fig 3a-c). We focus on how an individual cell’s state changes as it experiences different, controlled levels of inhibition from two immobilized and de-coupled neighbour cells. Lateral inhibition from these two neighbours (“External Dll4” or *D*_*ext*_) affects the individual EC under investigation (its current Dll4 level, “*D*_cell_”, is used as proxy for its active /inhibited state), but these neighbours are uncoupled from being inhibited themselves (Fig 4a). *D*_*cell*_is therefore analogous to the “ball position” in the mechanical example of a bistable system described in the results section and the fixed neighbours *D*_*ext*_ to the magnet position; raising *D*_*ext*_ can push the cell into a more inhibited (low *D*_cell_) state and vice versa.

In a hysteresis analysis, the external driving signal is quasi-statically raised and lowered in such a way that the dynamical variable stays at or close to its equilibrium value as the driving signal is varied. We start with *D*_*ext*_ set to zero and thus simulate a fully active EC with long filopodia (Fig. 3c). We then slowly raise the *D*_*ext*_ in increments of 50 until full inhibition is achieved (Fig. 3b), then reverse the process until it is yet again relieved. If filopodial active perception indeed creates bistability, we expect the transition between active /inhibited states to take place abruptly, at a *D*_*ext*_ threshold that is higher for the active →inhibited transition than the reverse.

##### Assessing quasi-static equilibrium

As the model is stochastic, we utilize a method to evaluate the quasistability of the system before incrementing the *D*_*ext*_ level as in hysteresis studies the system is expected to reach steady state before incrementing. In order to approximate quasi-static equilibrium while changing *D*_*ext*_, we tested the stability of *D*_*cell*_levels using the following method: 1) we allowed a transient time window (250 time-steps) to pass, 2) we calculated the average and standard deviation of *D*_*cell*_in two adjacent, non-overlapping sliding windows of size 250, 3) we tested whether 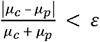 and 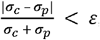, where 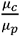 denote the average *D*_*cell*_and 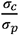 its standard deviation in the last /previous window (ε = 0.5). If the EC does not pass the test or a maximum time limit 60,000 was not yet reached, we increment the window. Otherwise we move on to the next *D*_*ext*_.

##### Characterizing *D*_*cell*_ fluctuations at intermediate levels in Calibrating quasi-static equilibrium assessment

We arrived at the generous • threshold value, used to assess the quasi-static equilibrium, by observing the fluctuations of *D*_*cell*_ close to the bistable region. Due to the stochastic nature of filopodia extensions, *D*_*cell*_ fluctuations close to the bistable region are large (Supp.Fig. 3a,b). Because of this inherent stochasticity, the exact *D*_*ext*_ thresholds at which abrupt transitions occur are not the same from simulation to simulation. Moreover, if the time window between increments is very large (well above the biological time scales involved in patterning), the EC may undergo stochastic transitions between metastable active /inhibited states. To understand the nature of these transitions, we set up a second set of simulations. In this case, we started the EC with *D*_*ext*_ =0 and ran the simulation for 10^4^ time steps, after which we abruptly increased D_*ext*_ to a nonzero target value, and performed 3· 10^4^ more steps. Exposure to low *D*_*ext*_ (500) keeps the EC in an active state, albeit with sizable *D*_*cell*_ fluctuations (Supp. Fig. 3c). At the intermediate *D*_*ext*_ level of 850, however, the long simulation showcases abrupt, stochastic transitions between active and inhibited states (Supp.Fig. 3d). Lastly, high *D*_*ext*_ (950) causes a one-way transition from active to a strongly inhibited state (Supp.Fig. 3e). Even here, the transition is not immediate (∼0.3 · 10^4^ steps), but it is irreversible within the time window of the run.

#### 3. Active Perception Signature Methods

##### The Model Setup

For work involving the AP signatures, we set up a minimal two cell MSM vessel.

##### Time to Pattern

Rather than the Vegfr and filopodia criteria as in the first MSM simulations, here we utilised the simpler and filopodia-independent *D*_*cell*_approach as developed in the hysteresis simulations above to determine the earliest time indicator of when the dynamical system has gone past the point of no return into the attractor of tip vs stalk. We simply measured the total Dll4 levels of the two cells and declared the system “Patterned” once one of the cells has maintained an advantage of 600 ligands over the other for 100 timesteps (the 100 timesteps of waiting were subtracted from the final number once elapsed). Once a cell achieves an advantage of 600 Dll4 (a value 6% of *D*_*max*_, decided on heuristically), in the absence of oscillation (which we test by waiting 100 time steps), the dynamics of the system seem to always cause that cell to be selected as the tip. This is a heuristic method and makes many assumptions such as the absence of a long-timescale oscillation and the presence of the bistable dynamics we witnessed in section 2 of this paper, reasonable for the constrained parameter ranges of F, V and G investigated here. The second assumption does not render the claims made in this paper circular, because the measures /heuristics used to characterize the bistability (thus ruling out oscillations) were different from those used that involved assuming its presence.

##### Sensory Map

To calculate the sensory map, we first calculated the displacement vector from the VEGFR activation occurring on memAgents existing away from the main cell body (e.g. on filopodia) to the centre of the circular cross section of the cylindrical cell body. Then we normalized this displacement vector and multiplied it by the radius of the blood vessel to point to the closest region on the surface of the cell to the activation. We mapped all activations out in space onto the cell surface in this way. One caveat of this approach is the activations from one cell can be attributed to the neighbour cell surface area if the filopodia reaches over it. However, the sensory map here represents the vessel level activations and sensing of the environment rather than precise individual cell sensing levels.

##### Estimating Entropy

To estimate Shannon entropy, we used the standard binning approach and Laplace smoothing to turn the data into a distribution. We did not have to quantize any data because of the discrete nature of Vegfr activations and the discrete spatial grid that the MSM model exists on.

##### Parameterisation

Parameters are detailed in Table 1.

**Table 1.**
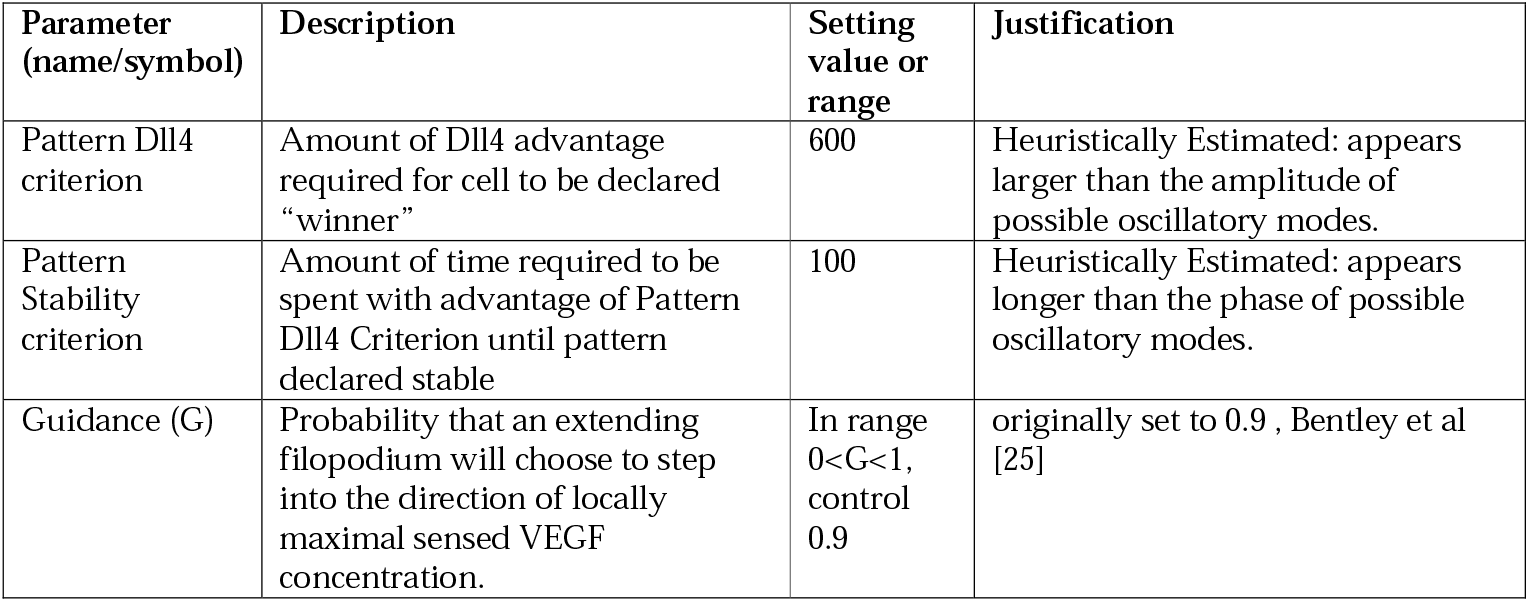
parameter table for in silico simulations.

##### Experimental Methods

###### Animals

Zebrafish embryos, larvae, and adults were maintained according to UK Home Office regulations. All studies were approved by the University of Manchester Ethical Review Board. Zebrafish strains were maintained at pH 7.4, a temperature of 28°C and exposed to 14 h light and 10 h dark cycles. Zebrafish lines used in this study were the Tg(kdrl:nlsEGFP)zf109 strain [100] and Tg(kdrl:ras-mCherry)s896 strain [101]

###### Time-lapse imaging

For confocal microscopy of live Tg(kdrl:nlsEGFP)zf109 and Tg(kdrl:ras-mCherry)s896 embryos was performed as previously described [20, 27]. Tg(kdrl:nlsEGFP)zf109 embryos were manually dechorionated and incubated with EtOH carrier or 0.08 µg /ml latrunculin B from 16 hpf prior to live imaging at 19 hpf. Tg(kdrl:ras-mCherry)s896 embryos were manually dechorionated prior to imaging at approximately 24 hpf. Embryos were mounted in 1% low-melt agarose in glass bottom dishes, which were subsequently filled with media supplemented with 0.0045% 1-Phenyl-2-thiourea and 0.1% tricaine, as well as EtOH carrier or 0.08 µg /ml latrunculin B for Tg(kdrl:nlsEGFP)zf109 embryos. Tg(kdrl:nlsEGFP)zf109 embryos were imaged using a 20x dipping objectives on a Zeiss LSM 700 confocal microscope. Tg(kdrl:ras-mCherry)s896 embryos were imaged using a 20x dipping objectives on a Zeiss LSM 880 confocal microscope with Airyscan. Embryos were maintained at 28C and stacks were recorded at every 0.3 h for Tg(kdrl:nlsEGFP)zf109 embryos or every 0.3 s for Tg(kdrl:ras-mCherry)s896 embryos. Tracking of cell motility was performed in ImageJ using the manual tracking plugin. All cell tracking recordings were normalized at each time point relative to the position of the dorsal aorta to account for any dorsal or ventral drift of embryos during imaging.

###### pErk immunostaining

Whole-mount immunostaining for pErk was performed as previously described [20, 27]. Briefly, Tg(kdrl:nlsEGFP)zf109 embryos were manually dechorionated and incubated with EtOH carrier or 0.08 µg / ml latrunculin B from 16-18 hpf prior to fixation in PFA at the indicated time points. Embryos were washed in MeOH, bleached in 3% H202 in MeOH, washed again in MeOH, stored in MeOH at −20°C for 2 days in MeOH before equilibration with PBT (PBS, 0.1% Tween-20). Embryos were cryoprotected in 30% sucrose in PBT before further equilibration in PBT, incubation with 150 mM Tris-HCl (pH 9.0) and heating to 70°C for 15 min. Embryos were then, washed in PBT, dH2O, acetone, PBT and TBST (TBS, 0.1% Tween-20, 0.1% Triton X-100), prior to incubation overnight at 4°C with block solution (TBST, 1% BSA, 10% goat serum). Embryos were subsequently incubated overnight with anti-phospho-ERK1 /2 antibody (1:250, Cell Signaling; #4370) in blocking buffer, washed with TBST, washed with Maleic buffer (150 mM Maleic acid, 100 mM NaCl, 0.001% Tween-20, pH 7.4), blocked in 2% blocking reagent (Sigma-Aldrich) in Maleic buffer for 3 h and incubated with goat anti-rabbit IgG-HRP (1:1000) in 2% blocking reagent in Maleic buffer overnight at 4°C. Embryos were then washed in Maleic buffer and then PBS at room temperature prior to incubation with 50 µl amplification diluent with 1 µl Tyramide-Cy3 (Perkin Elmer) for 3 h at room temperature in the dark. Embryos were finally washed over several days in TBST at room temperature. Embryos were then mounted in 1% low-melt agarose in glass bottom dishes and imaged using a 20x dipping objectives on a Zeiss LSM 700 confocal microscope. Finally, to identify pErk-positive endothelial cells, kdrl:nlsEGFP signal was used to create a binary channel mask to isolate only nuclear signal in the pErk channel.

## Acknowledgments

We would like to thank Dylan Feldner-Busztin for Python data science support. K.B, B.Z, T.M, K.V-V, I.A were supported by The Francis Crick Institute, which receives its core funding from Cancer Research UK (FC001751), the UK Medical Research Council (FC001751), and the Wellcome Trust (FC001751).S.P.H. was funded by the Wellcome Trust (Ref. 219500 /Z /19 /Z). S.P.H. and R.T. were funded by the British Heart Foundation (PG /18 / 67 /33891). G. C. is a BBSRC DTP PhD student. K.B and E.R.R were funded in part by BIDMC for early work on the project while located at BIDMC. E.R.R was funded by BIDMC and NIH grant HL077348-03 while at BIDMC during early work towards this study, then by the College of Wooster (Henry Luce III Funds) and NIH grant HL119322-04. K.I.H was supported by institutional training grant T32 HL07893 from the NHLBI of the NIH during early work on this project while located at BIDMC.

---

**Supplemental Figure 1.**
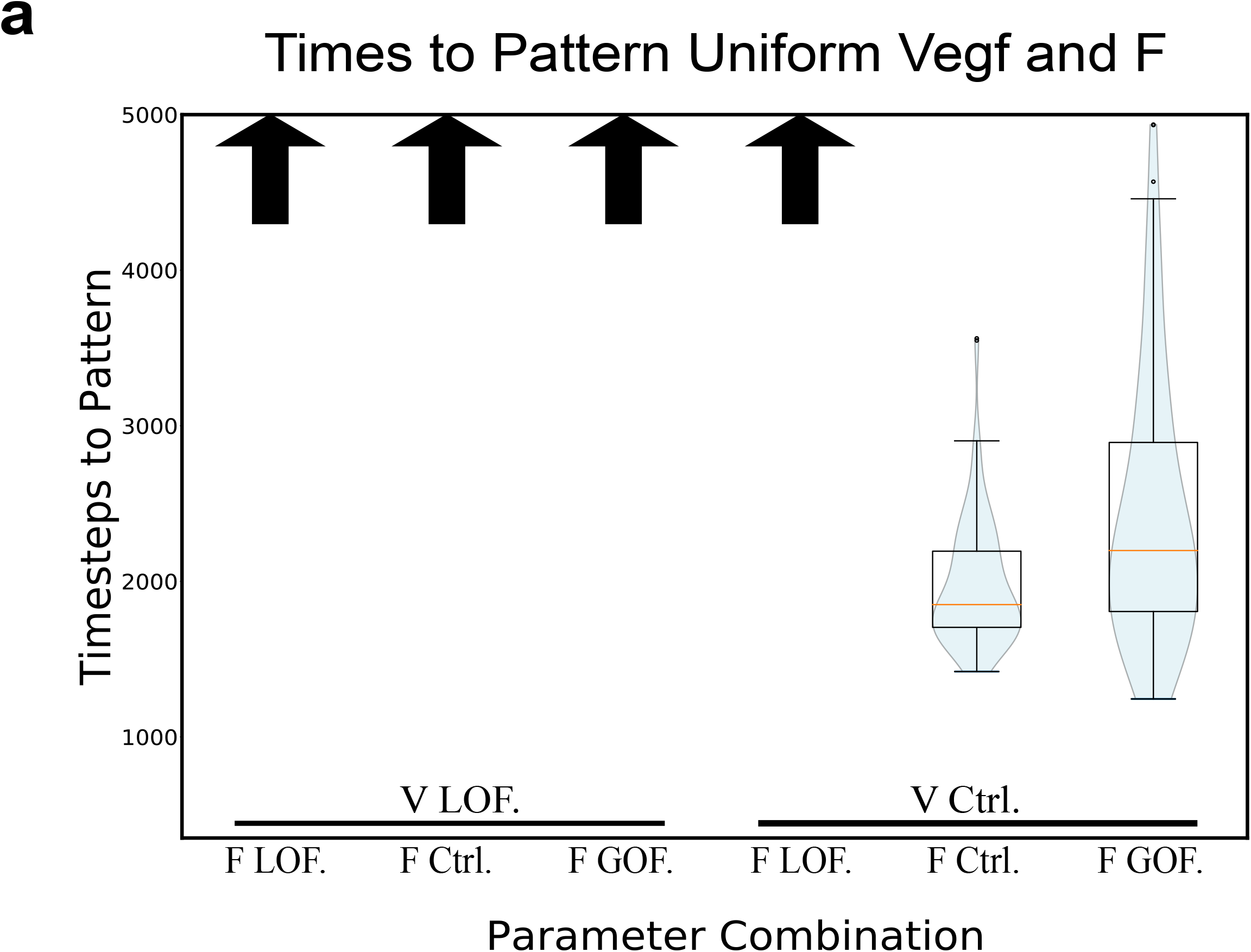
Loss of Gradient slows down Notch selection. **a)**Violin plots showing that in completely uniform Vegf (no gradient) slows down selection (“time to pattern”) of the *in silico* for control (0.8) and LOF Vegf (0.4) concentrations compared to simulations with a vegf gradient (Fig 2b,c). Altering filopodial activity fails to rescue this. Blue shading corresponds to estimated probability density of timing values in that range. Upward arrows indicate pattern not achieved within the time range of the simulation for the corresponding parameter settings. Horizontal pink line indicates median value, horizontal black lines indicate middle quartiles, whiskers indicate upper and lower quartiles. N=100 simulations ran for 5000 timesteps.

**Supplemental Figure 2.**
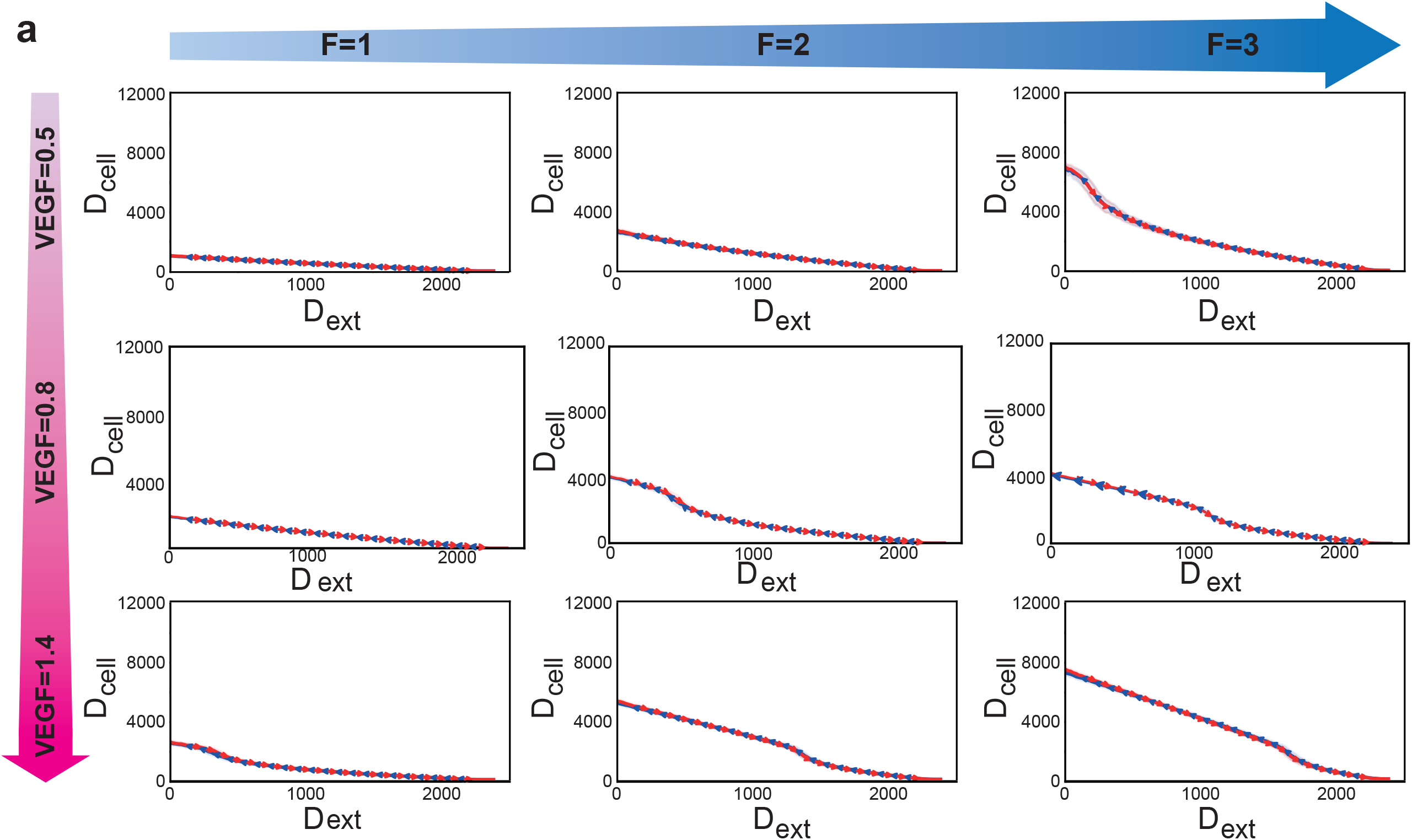
Vegf gradient required for bistable Dll4 dynamics. **a)** Hysteresis analysis for uniformly distributed environmental Vegf (no gradient) covaried with Filopodial activity (F) shows that the bistability and hysteresis is lost upon loss of VEGF gradient, regardless of the concentration of Vegf present in environment. This shows the dependence of the bistable dynamics on Vegf-based positive feedback. N=100, shading indicates 1 standard deviation.

**Supplemental Figure 3.**
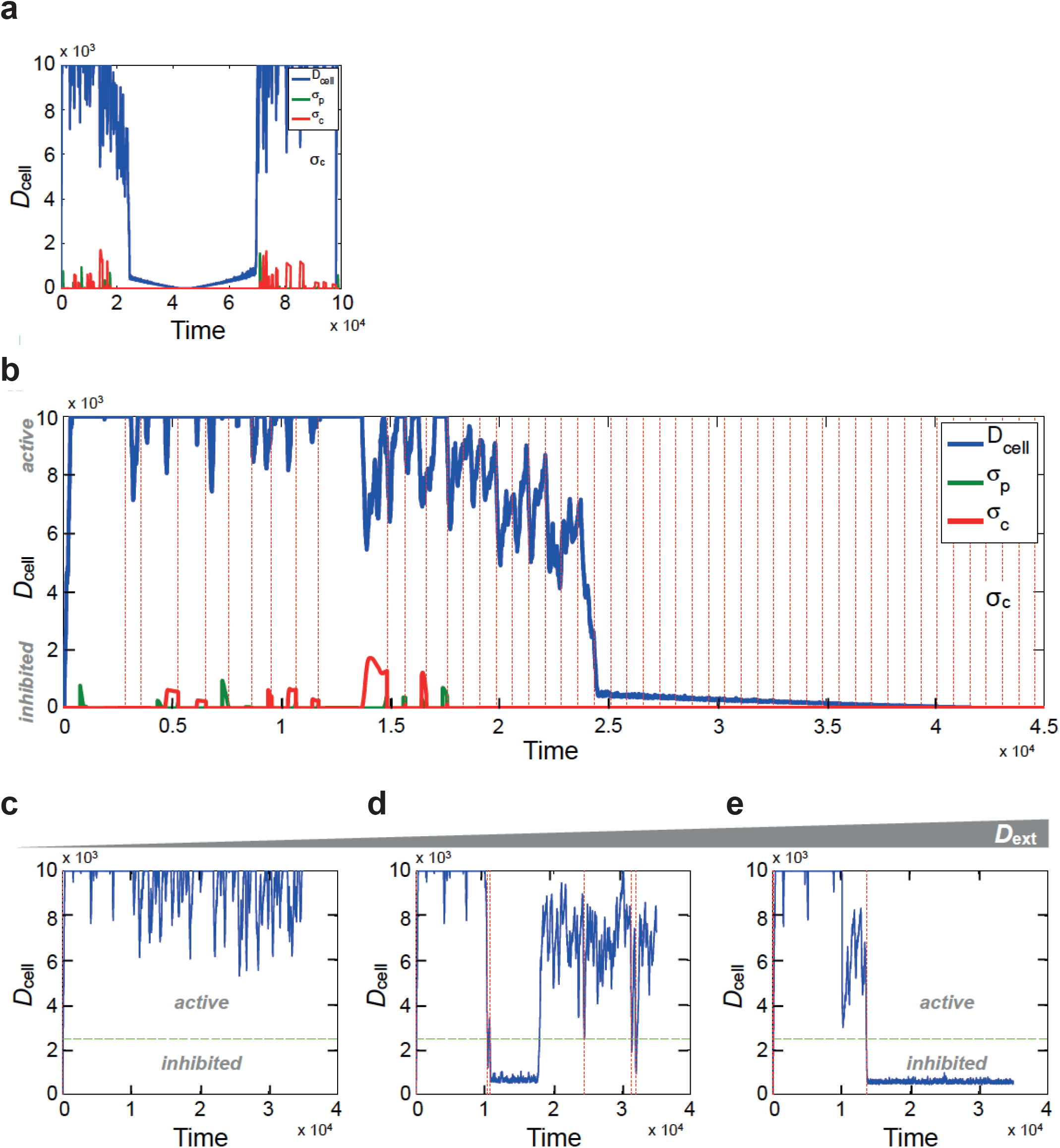
Hysteresis Details. **a)** Time course of *D*_*cell*_during a full cycle of hysteresis (blue). Standard deviation of *D*_cell_, measured in two non-overlapping, expanding windows (*σ*_*p*_ for previous and *σ*_*c*_ for current windows) are shown in green and red, respectively. **b)** Zoomed-in time course of *D*_*cell*_during a half-cycle of hysteresis, showing large *D*_*cell*_fluctuations in the active state as the cell re-stabilises on each *D*_*ext*_ increment. Red lines: times of *D*_*ext*_ increment). **c**,**d**,**e)** Long *D*_*cell*_time-courses of ECs exposed to fixed *D*_*ext*_. Green line: rarest observed *D*_*cell*_value, marking the position of the barrier between tip and stalk states (see Fig 3e). Red line: transitions between active / inhibited states. Left: *D*_*ext*_ = 500; EC restricted to the active state. Middle: *D*_*ext*_ = 850; bistable EC stochastically transitioning between active and inhibited states. Right: *D*_*ext*_ = 950; EC restricted to its inhibited state.

